# Diagnosing protein sequence search in the era of language models

**DOI:** 10.64898/2026.04.26.720921

**Authors:** Han Zhou, Yifan Yang, Yang Young Lu

## Abstract

Protein language model (PLM) based search is rapidly emerging as a successor to classical sequence alignment, with recent high-profile studies reporting substantial improvements in speed and remote homology detection. However, success on standard benchmarks does not guarantee that similarity derived from PLM embeddings constitutes reliable biological evidence. Here, we introduce PLM-GUARD, a diagnostic framework designed to interrogate the underlying meaning of protein search scores and assess their biological trustworthiness. PLM-GUARD comprises six sanity checks spanning biological fidelity, semantic validity, and manipulation safety. Across eight representative search methods, classical alignment-based systems demonstrate remarkable robustness, whereas current PLM-based methods fail broadly across all three dimensions. Notably, hybrid methods show intermediate results, indicating that alignment is still critical for ensuring biologically grounded correspondence. Our findings provide a timely clarification for the field and underscore the necessity of diagnostic evaluation as protein search enters the era of language models.

## 1 Introduction

Protein sequence search is a foundational operation in biological research, supporting tasks that range from homology detection and functional annotation to evolutionary analysis and structure prediction [1–4]. For decades, alignment-based methods have dominated this space [5, 6], in part because their scoring functions display predictable and biologically grounded behavior. Alignment scores typically decrease as sequences diverge through evolution, increase as additional matching residues are aligned, and can be interpreted through explicit substitution models and gap penalties [7, 8]. Because of these properties, alignment-based similarity has served as more than a retrieval criterion. In practice, it has also functioned as meaningful evidence for downstream biological inference [9].

Protein language models (PLMs) have introduced a qualitatively different paradigm for sequence search [10]. By mapping sequences into dense learned representations, these models enable similarity assessment without explicit residue-level alignment (Figure 1a) [11]. This shift has improved computational efficiency and has shown strong retrieval performance across many benchmarks, particularly in the so-called “twilight zone” of remote homology [12]. As a result, PLM-based protein sequence search has been rapidly adopted, with several recent studies presenting PLM similarity as a powerful alternative to traditional alignment for large-scale search and remote homology detection [13–16].

**Fig. 1:**
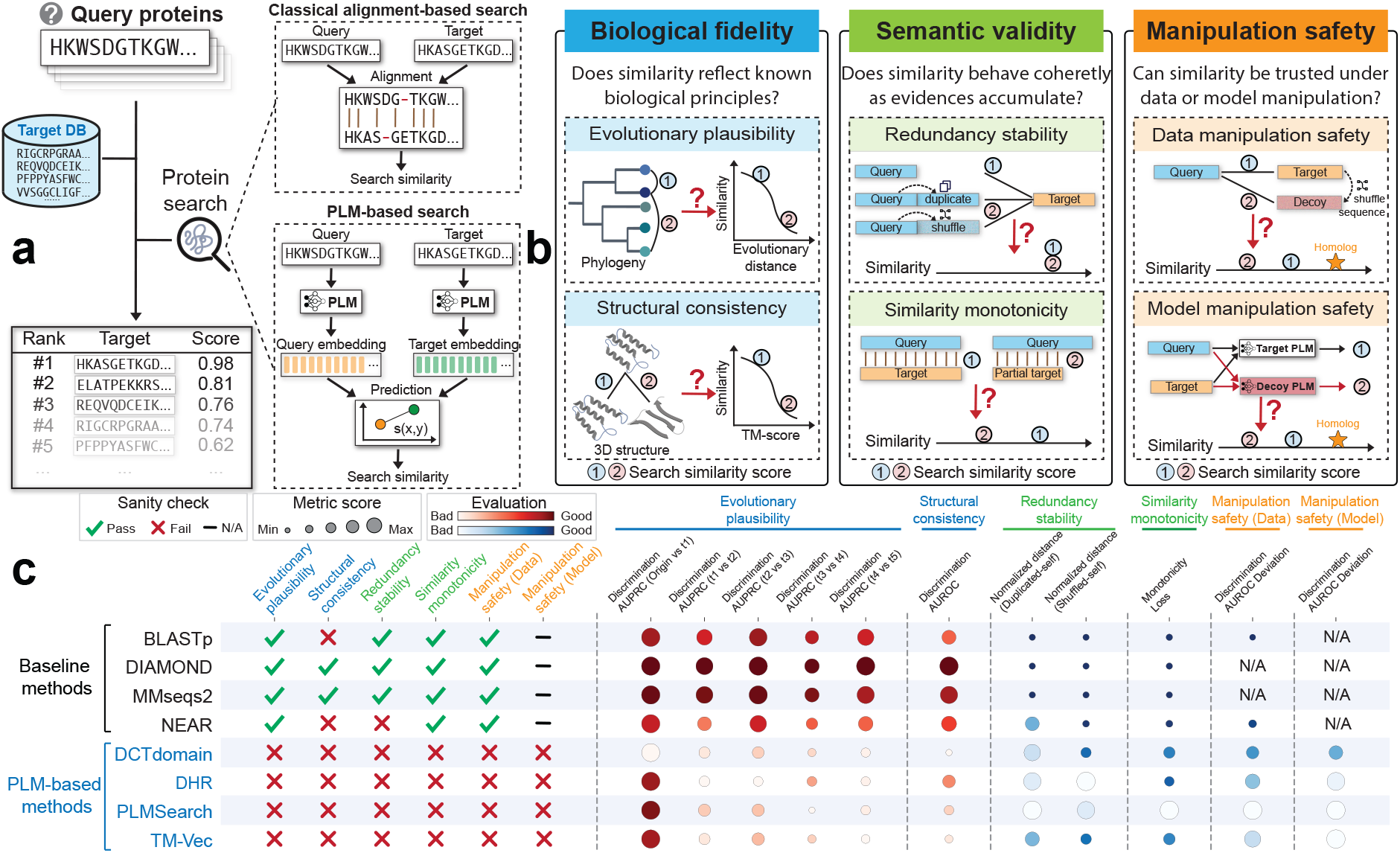
Overview and sanity check summary of PLM-GUARD. **(a)** Classical alignment-based and PLM-based search produce similarity through fundamentally different mechanisms. **(b)** PLM-GUARD organizes six sanity checks into three dimensions: biological fidelity, semantic validity, and manipulation safety. **(c)** The summary across eight representative methods shows pass or fail outcomes for each sanity check, with circles indicating the corresponding quantitative scores.

Despite this progress, a fundamental question remains insufficiently examined: “what does a PLM-based similarity score actually represent?” Benchmark metrics can show that a method retrieves biologically related sequences, but they reveal much less about the meaning and reliability of the similarity scores themselves [17]. Unlike alignment scores, which are constrained by explicit biological assumptions, PLM-based similarities arise from learned representations whose underlying constraints are implicit and often difficult to interpret [18]. Existing evaluations rarely test whether these scores track evolutionary distance in a biologically sensible way, behave predictably as matching evidence is added or removed, or resist spurious inflation in the absence of genuine biological relatedness [11, 19]. More broadly, the usefulness of sequence similarity as evidence depends not only on retrieval success, but also on whether the connection between score and biological meaning has been explicitly examined.

These considerations motivate a diagnostic view of protein sequence search in the era of language models. Rather than asking which method performs best under a fixed benchmark [20], we ask whether similarity scores satisfy a set of basic properties required for them to function as meaningful and trustworthy inferential signals [21]. To this end, we introduce PLM-GUARD, a diagnostic framework built around six sanity checks organized into three dimensions: biological fidelity, semantic validity, and manipulation safety (Figure 1b). Biological fidelity asks whether similarity scores remain consistent with core biological relationships, including evolutionary divergence and structural conservation. Semantic validity asks whether scores behave coherently when query content is altered in controlled ways, remaining stable under redundancy while changing predictably as informative sequence content is removed. Manipulation safety asks whether scores resist artificial inflation, both when the data are perturbed synthetically and when the representation pipeline itself is manipulated. Together, these tests provide a systematic way to examine how PLM-based similarity behaves across biologically and methodologically relevant scenarios. Unfortunately, the PLM-based search methods examined in this study exhibit broad failures across these sanity checks (Figure 1c), indicating that high embedding similarity may not reliably serve as biological evidence without further validation. Our goal is not to dispute reported gains in retrieval performance, but to emphasize the need for a more cautious and diagnostically informed use of PLM-based search.

## 2. Results

### 2.1 PLM-GUARD defines six sanity checks for protein search similarity

PLM-based sequence search replaces explicit alignment with proximity in representation space, yet the resulting similarity scores are often interpreted as if they were direct biological evidence. This shift makes it necessary to examine not only whether a method retrieves related sequences, but also whether its similarity scores behave in ways that support meaningful interpretation. To address this question, our diagnostic framework instantiates the three dimensions introduced above through six concrete sanity checks, each designed to probe a distinct aspect of similarity behavior. Collectively, these checks define a set of conditions under which PLM-based similarity can be interpreted as meaningful evidence rather than as an opaque retrieval signal.

The first dimension, biological fidelity, asks whether similarity remains consistent with basic biological relationships. It is examined through the evolutionary plausibility and structural consistency tests. The evolutionary plausibility test evaluates whether similarity decreases in a manner consistent with evolutionary divergence [7]. The structural consistency test asks whether similarity tracks three-dimensional relatedness even when primary sequence identity has largely disappeared [22]. A method that satisfies biological fidelity should assign lower similarity scores as sequences diverge, yet still recognize structurally related proteins in the remote-homology “twilight zone”, even when controlled perturbations weaken sequence identity while preserving structure [23].

The second dimension, semantic validity, asks whether similarity behaves coherently as sequence evidence is altered in controlled ways. This dimension is evaluated through the redundancy stability and similarity monotonicity tests. Redundancy stability examines whether similarity remains stable when redundant or irrelevant query content is appended. Similarity monotonicity examines whether scores change predictably as informative sequence content is progressively reduced or restored [24]. A method that satisfies semantic validity should not reward redundancy, but should respond coherently when the amount of meaningful evidence changes.

The third dimension, manipulation safety, asks whether similarity can be artificially inflated without genuine homology. It is evaluated through the data manipulation safety and model manipulation safety tests. Data manipulation safety measures the susceptibility of similarity scores to spurious inflation by synthetic null sequences that lack true homology [25], including decoys constructed by shuffling target sequences in the database. Model manipulation safety examines whether similarity remains stable when PLM-derived representations are perturbed in a controlled way [26]. A method that supports trustworthy interpretation should suppress similarity to such biological decoys and should not be easily redirected by modest perturbations of the representation pipeline.

These six sanity checks are intentionally model-agnostic and together operationalize the three diagnostic dimensions. Although similarity is measured on a continuous scale, each diagnostic tests whether a qualitative interpretability constraint is respected or violated. Failure on a given check therefore reflects a breakdown in interpretability under the corresponding condition. To ground these diagnostics, we evaluated a deliberately diverse set of search methods, including three classical alignment-based methods, four PLM-based methods, and one hybrid method that combines embeddings with alignment, as described in Section 4.1. The generally stable behavior of the alignment-based controls shows that the proposed tests are reasonable and practically satisfiable, rather than artificially stringent. The inclusion of the hybrid method further helps distinguish failures that are specific to fully embedding-based search from behaviors that are preserved when alignment remains part of the scoring procedure. Against this backdrop, the broad failures observed in PLM-based methods become more informative, because they point to specific breakdowns in how similarity is represented and interpreted.

### 2.2 PLM-based methods fail to track evolutionary divergence under controlled mutation

We first asked whether protein search similarity decays in a biologically coherent way under controlled evolutionary divergence. Following the protocol in Section 4.2, we simulated mutation trajectories from ASTRAL40 domains [27] and tracked the score assigned to the true homolog as divergence increased from the original sequence to progressively mutated descendants (Figure 2a). This setting provides a direct test of whether a method preserves the expected ordering of similarity under evolution. Because evolutionary plausibility is a basic requirement for treating similarity as evidence of relatedness, violations of this ordering indicate not just reduced utility, but a breakdown in biological interpretability.

**Fig. 2:**
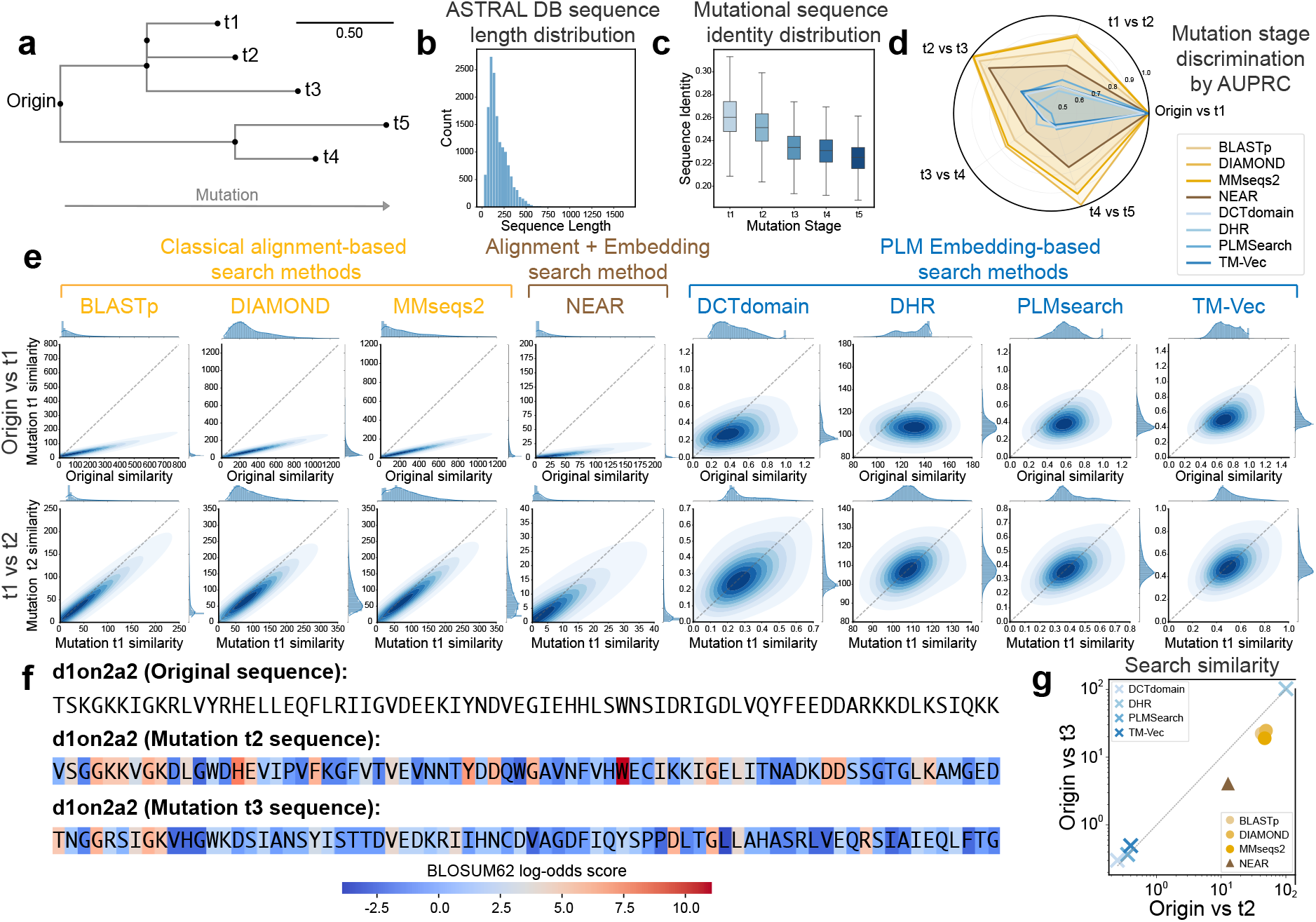
Evolutionary plausibility sanity check. Evolutionary plausibility tests whether protein search similarity decreases coherently under controlled sequence divergence. (**a**) Proteins from the ASTRAL40 database are evolved into progressively diverged descendants (*t*1–*t*5). (**b**) The ASTRAL40 database exhibits broad sequence-length variation for the evolutionary plausibility test. (**c**) Sequence identity decreases progressively across mutation stages and confirms controlled sequence divergence. (**d**,**e**) PLM-based search methods struggle to discriminate between adjacent stages of divergence, as reflected quantitatively and qualitatively. (**f**) Example original and mutated sequences, with substitutions colored by BLOSUM62 log-odds score. (**g**) Continuation of the example in (f), showing similarity scores for the same query-target pair under increasing divergence (Origin-t2 vs. Origin-t3). Non-PLM methods preserve the expected monotonic decrease in similarity, whereas PLM-based methods often violate this monotonic behavior.

The simulation setup produced a heterogeneous and well-controlled evaluation background. The ASTRAL40 database spans a broad range of sequence lengths (Figure 2b), and sequence identity to the original sequence decreases progressively across mutation stages (Figure 2c), confirming that the synthetic trajectories capture controlled divergence rather than arbitrary perturbation. Under this setup, methods with strong evolutionary plausibility should assign systematically lower similarity to increasingly diverged variants and should discriminate adjacent mutation stages with high consistency.

This expectation was largely met by alignment-based methods and, to a lesser extent, by NEAR. Classical alignment-based methods, including BLASTp, DIAMOND, and MMseqs2, effectively distinguished between adjacent mutation stages. NEAR remained substantially better behaved than fully PLM-based methods, indicating that the retention of alignment helps maintain biologically coherent score decay even when embeddings are incorporated. In comparison, PLM-based methods, including DCTdomain, DHR, PLMSearch, and TM-Vec, exhibited substantially weaker discrimination between adjacent mutation stages, as reflected by lower AUPRC values (Figure 2d) and the corresponding qualitative analysis (Figure 2e). Although the mutation stages differ in their degree of divergence, these methods consistently struggle to separate neighboring stages across the trajectory (Figure S1).

Lastly, the failure mode is illustrated at the level of individual examples (Figure 2f). In the comparison between Origin–*t*2 and Origin–*t*3, alignment-based methods generally preserve the correct similarity ordering, assigning higher similarity to the less diverged sequence and lower similarity to the more diverged one (Figure 2g). PLM-based methods more often compress these differences or fail to maintain the expected ordering, even though the mutational process itself remains well controlled. Taken together, these results show that current PLM-based search methods are substantially less reliable at preserving evolutionary plausibility, which limits the extent to which their similarity scores can be interpreted as direct evidence of evolutionary relatedness.

### 2.3 PLM-based methods fail to preserve structural consistency under structure-preserving mutation

We next asked whether protein search similarity remains consistent with three-dimensional structural relatedness, especially in the remote-homology regime where sequence identity alone provides limited guidance. Following the protocol in Section 4.3, we generated structure-preserving mutated queries by introducing sequence perturbations and then relaxing the resulting structures (Figure 3a). This design allows us to test whether a method continues to recognize structurally related homologs even after the query sequence has been altered, while also distinguishing them from non-homolog controls. Because structural conservation often persists after obvious sequence similarity has weakened, failure in this setting indicates that similarity is not reliably tracking biologically meaningful structural relationships.

**Fig. 3:**
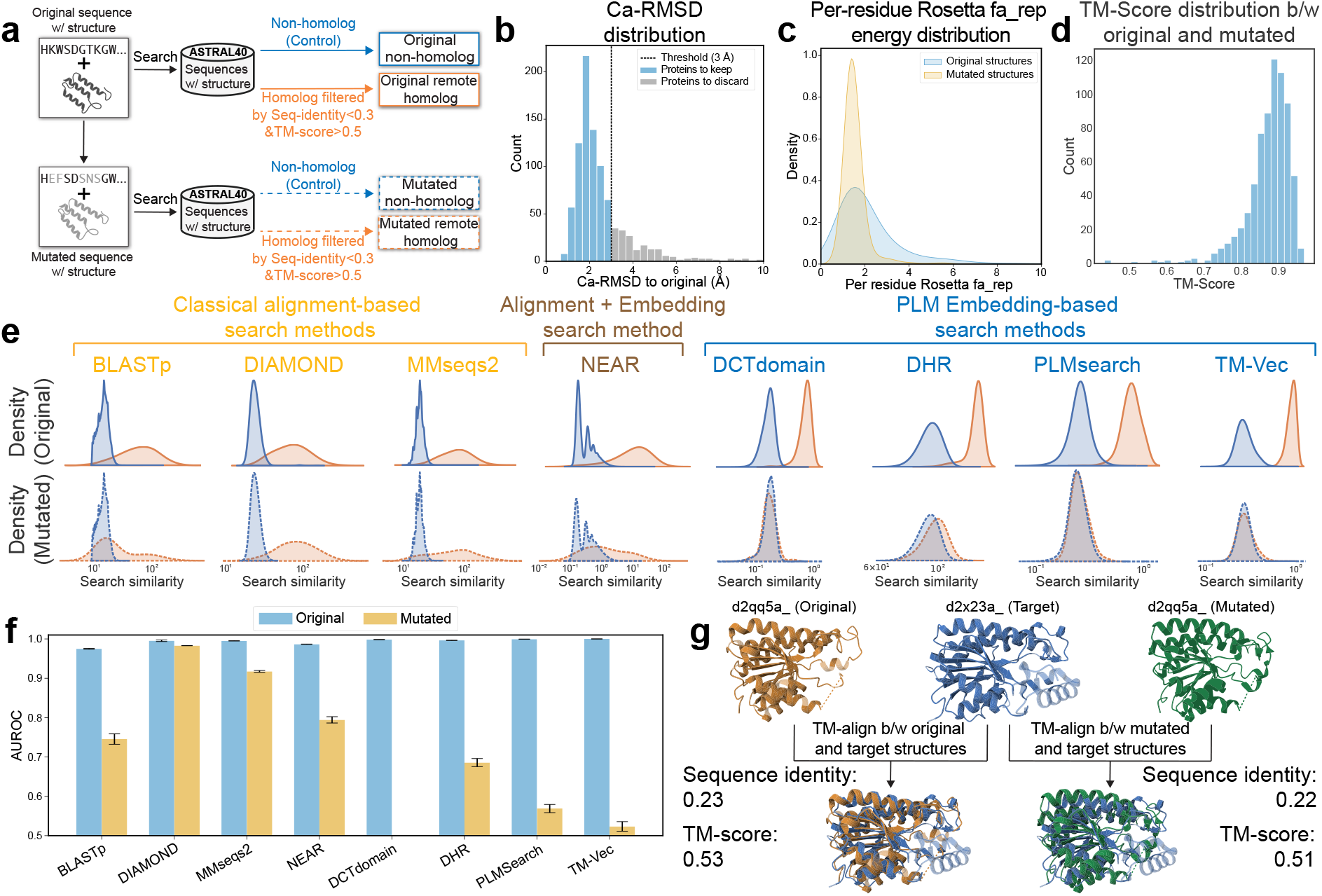
Structural consistency sanity check. Structural consistency tests whether protein search similarity remains consistent with structural relatedness after controlled sequence mutation followed by structural relaxation. (**a**) The structural consistency test compares retrieval behavior for original and mutated queries against remote homologs and non-homolog controls drawn from ASTRAL40 structures. (**b**,**c**) The relaxed mutated structures remain physically reasonable, with most mutated structures staying close to their original structures in C*α*-RMSD and showing similar per-residue Rosetta fa rep energy distributions. (**d**) TM-scores between original and mutated structures remain high, confirming substantial structural similarity despite sequence perturbation. (**e**) PLM-based methods show large distributional shifts after mutation, indicating weak structural consistency, whereas non-PLM-based methods remain comparatively stable. (**f**) PLM-based search methods struggle to discriminate remote homologs from non-homologs under structure-preserving mutation, as reflected quantitatively by AUROC. (**g**) An example remote homolog pair shows that mutation alters sequence identity while leaving the TM-score nearly unchanged, illustrating that the structural relationship is preserved despite sequence perturbation.

The mutated structures remained suitable for this evaluation. Most stayed close to their original structures in C*α*-RMSD and exhibited similar per-residue Rosetta fa rep energy distributions, indicating that the relaxation procedure did not produce unrealistic conformations (Figure 3b,c). TM-scores between original and mutated structures also remained high (Figure 3d), confirming that the overall structure was largely preserved despite substantial sequence perturbation. Example remote homolog pairs further illustrate this point: mutation changes sequence identity while leaving the TM-score nearly unchanged, showing that the structural relationship is retained even when sequence similarity is weakened (Figure 3g and Figure S1).

Under this structure-preserving perturbation, non-PLM-based methods remained comparatively stable, whereas PLM-based methods showed much larger shifts in their search-similarity distributions (Figure 3e). DIAMOND and MMseqs2 retained the clearest structural consistency, and NEAR again occupied an intermediate position, suggesting that alignment continues to play an important role in anchoring similarity to biologically meaningful correspondence. BLASTp was less robust than DIAMOND and MMseqs2, consistent with its weaker behavior in the remote-homology setting. This pattern is also reflected in discriminating remote homologs from non-homolog controls. PLM-based search methods struggled more under structure-preserving mutation, as quantified by AUROC (the area under a receiver operating characteristic curve) (Figure 3f). In other words, even when the structural relationship remains largely intact, their similarity scores become much less reliable at separating true remote homologs from unrelated proteins. Taken together, these results indicate that current PLM-based search methods do not consistently preserve structural consistency, which limits the extent to which their similarity scores can be interpreted as evidence of structural relatedness in low-identity regimes.

### 2.4 PLM-based methods fail redundancy stability under non-informative query augmentation

We next evaluated whether protein search similarity remains stable when redundant or non-informative sequence content is added to the query. Following the protocol in Section 4.4, we compared homolog similarity scores from the original query with scores obtained after appending either an exact duplicate of the query or a shuffled version with the same composition (Figure 4a). Under this diagnostic, a method satisfies redundancy stability only if these perturbations leave similarity to true homologs largely unchanged. If scores increase simply because the query becomes longer or more repetitive, then similarity is no longer tracking biological evidence in a coherent way.

**Fig. 4:**
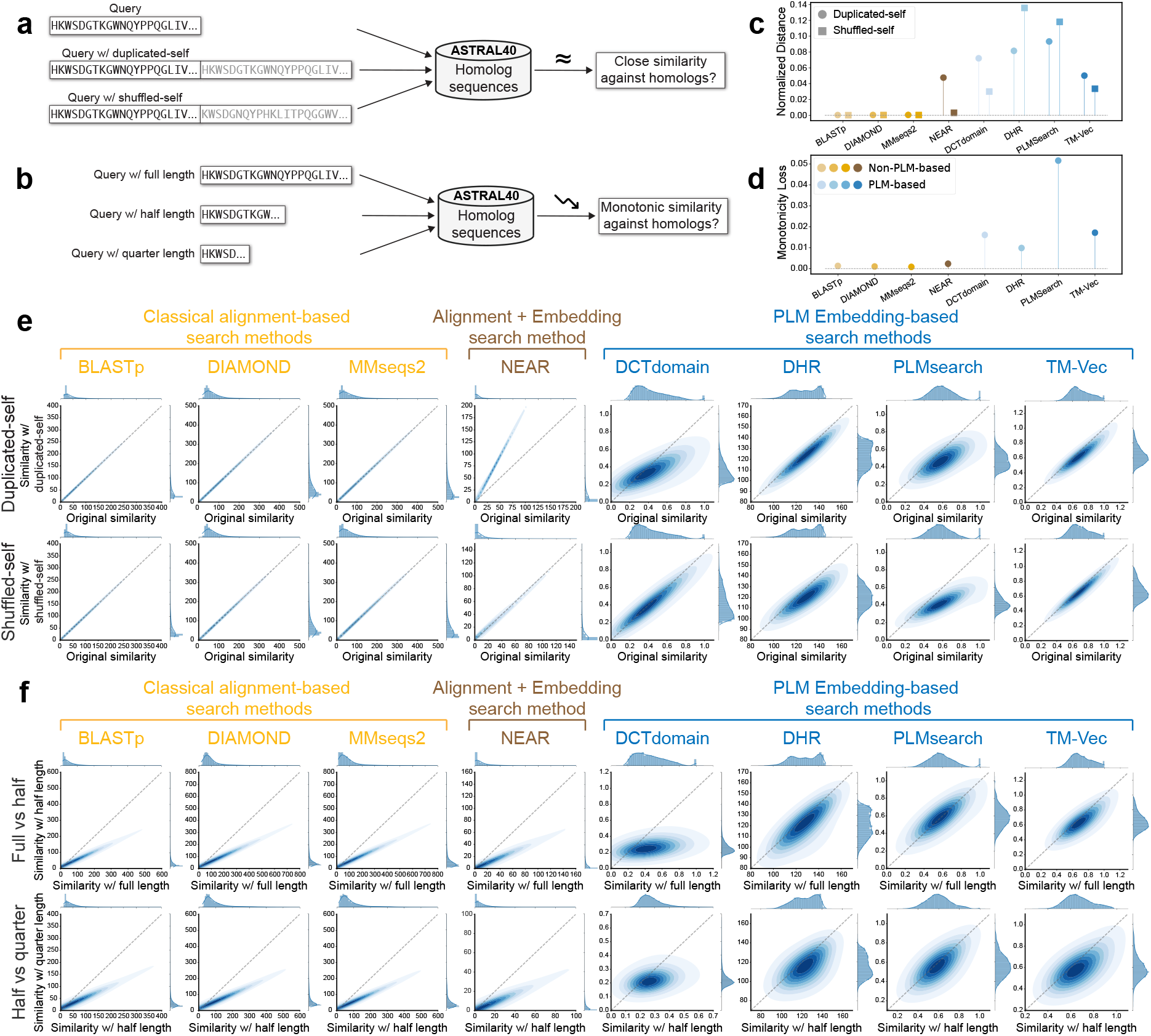
Semantic validity sanity check. Semantic validity tests whether protein search similarity behaves coherently when sequence evidence is redundant or partially removed. (**a**) The redundancy stability test evaluates whether duplicating or shuffling the query preserves similarity against homologs in ASTRAL40. (**b**) The similarity monotonicity test evaluates whether similarity to homologs decreases predictably as the query is progressively truncated. (**c**) PLM-based methods exhibit redundancy instability under redundant or non-informative sequence perturbations, as reflected by large normalized distances. (**d**) PLM-based methods fail to preserve similarity monotonicity under progressive query truncation, as reflected by large monotonicity loss, whereas non-PLM-based methods remain largely monotonic. (**e**,**f**) PLM-based methods exhibit redundancy instability under duplicated-self and shuffled-self perturbations and fail to preserve the expected monotonic decrease in similarity under query truncation, whereas non-PLM-based methods remain comparatively stable.

This diagnostic quantitatively reveals a clear difference between non-PLM-based and PLM-based methods. Classical alignment-based methods maintain stability when duplicate or shuffled sequences are appended to the query, suggesting that redundant or non-informative content does not substantially distort similarity assignments. In comparison, PLM-based methods show much larger deviations from the expected invariant behavior, as quantified by the normalized distance metric (Figure 4c). These results indicate that PLM-based similarity is often sensitive to query augmentation even when no new biological information is introduced.

Qualitatively, non-PLM-based methods remain closer to the expected diagonal relationship between original and variation-query similarity, whereas PLM-based methods show more pronounced departures under both duplicated-self and shuffled-self perturbations (Figure 4e). NEAR behaves differently from both groups when the query is duplicated, because its similarity is computed by matching each query residue embedding to its best target residue embedding and then summing over all matched residue pairs. As a result, duplicating the query effectively doubles the residue-level evidence presented to the scoring function and therefore approximately doubles the overall score, even though no new biological information has been introduced. Under the diagnostic definition used here, this behavior constitutes a redundancy stability violation, because similarity becomes sensitive to repeated evidence rather than remaining invariant to it. Taken together, these results show that many PLM-based search methods violate redundancy stability, meaning that their similarity scores can be inflated by repeated or non-informative sequence content rather than by additional biological evidence.

### 2.5 PLM-based methods fail to preserve similarity monotonicity under controlled evidence removal

We next evaluated whether protein search similarity decreases coherently as informative sequence content is progressively removed from the query. Following the protocol in Section 4.5, we compared homolog similarity scores obtained from the full-length query with those obtained after truncating the query to shorter fractions (Figure 4b). Under this diagnostic, a method satisfies similarity monotonicity if reducing the amount of matching evidence leads to a corresponding decrease in similarity. If scores fail to attenuate, or do so irregularly, then similarity can no longer be interpreted as a coherent measure of evidence strength.

This diagnostic separates non-PLM-based and PLM-based methods. Non-PLM-based methods remain largely monotonic under progressive truncation, indicating that their scores respond in the expected direction as informative sequence content is removed. In comparison, PLM-based methods show substantially larger monotonicity loss (Figure 4d), defined as the failure for cases in which similarity does not decrease, or even increases, after informative sequence content is removed. In other words, truncating the query does not reliably weaken similarity in proportion to the loss of evidence.

The qualitative divergence between non-PLM and PLM-based methods is even more pronounced. For non-PLM-based methods, homolog similarity under truncation remains relatively well ordered with respect to the full-length query, whereas PLM-based methods together with NEAR show much less consistent structure and more frequent departures from the expected monotone relationship (Figure 4f). Although NEAR incorporates embeddings, it largely preserves similarity monotonicity, in contrast to fully PLM-based methods. This suggests that retaining alignment helps maintain a coherent relationship between the amount of matching evidence and the resulting similarity score. Taken together, these results show that many PLM-based search methods fail to preserve similarity monotonicity under controlled evidence removal. As a result, their scores are less reliable as quantitative indicators of how much biologically meaningful sequence evidence actually supports a match.

### 2.6 PLM-based methods are vulnerable to data-level hijacking by shuffled decoys

We next asked whether high similarity can be induced without genuine homology through direct manipulation of the input data. Following the protocol in Section 4.6, we augmented the target database with shuffled decoy sequences that preserve low-level composition while destroying biological meaning (Figure 5a). Under this diagnostic, a method satisfies data manipulation safety only if such decoys remain clearly separated from true homologs and behave similarly to ordinary non-homologs. If shuffled decoys can intrude into the high-similarity regime, then the resulting scores are vulnerable to artificial inflation rather than being anchored to genuine biological relatedness.

**Fig. 5:**
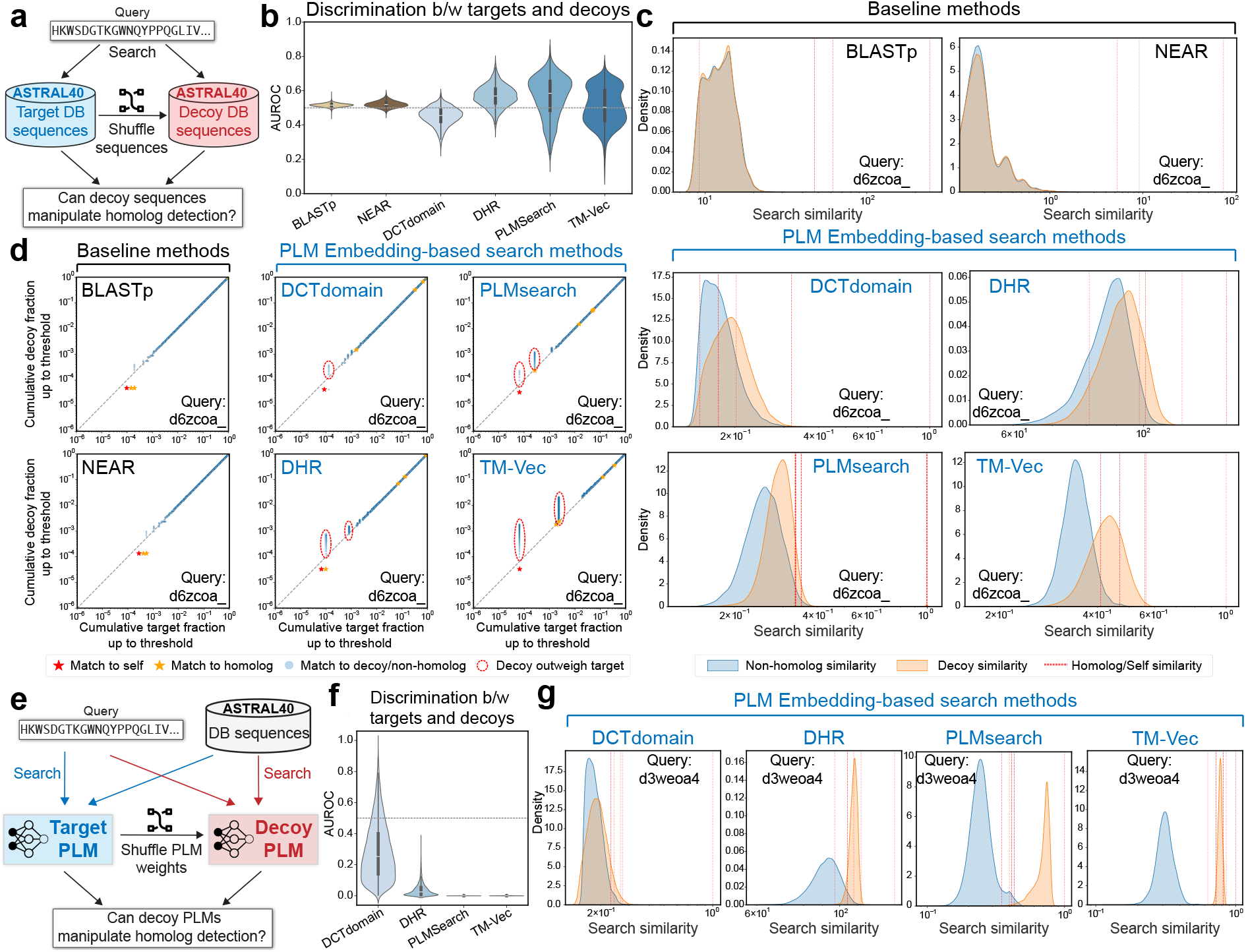
Manipulation safety sanity check. Manipulation safety tests whether high similarity can be hijacked without genuine homology through manipulation of the data or the model. (**a**) The data manipulation safety test asks whether shuffled decoy sequences can intrude into homolog detection. (**b**) PLM-based methods rank shuffled decoys much higher relative to targets, as reflected by more dispersed AUROC than BLASTp and NEAR; MMseqs2 and DIAMOND are omitted because their pre-filtering design filters out decoys before explicit similarity scores can be assigned. (**c**) An example query illustrates manipulation safety in non-PLM-based methods, where decoy similarities remain close to non-homolog similarities. (**d**) Cumulative target-versus-decoy curves show that PLM-based methods more readily admit thresholds where decoys outweigh targets, with such samples highlighted. (**e**) The model manipulation safety test asks whether a decoy PLM with shuffled weights can hijack homolog detection. (**f**) PLM-based methods rank decoy PLMs much more competitively against targets, as reflected by deviated AUROC under model manipulation; this applies only to PLM-based methods, not to classical search methods. (**g**) Example queries show that decoy PLMs can assign decoy or non-homolog matches similarities that exceed homolog matches, illustrating vulnerability to model-level hijacking.

This diagnostic reveals a sharp contrast between non-PLM-based and PLM-based methods (Figure 5b). BLASTp and NEAR show weak separation between targets and shuffled decoys, as most targets are non-homologs and should therefore be nearly as uninformative as shuffled decoys, a pattern reflected quantitatively by AUROC. MMseqs2 and DIAMOND are not included in this comparison because their pre-filtering strategy filters out decoys before explicit similarity scores can be assigned. In comparison, PLM-based methods show more variable separation between targets and shuffled decoys. Some methods, such as DCTdomain, rank decoys much more competitively against true targets, whereas others, including DHR, PLMSearch, and TM-Vec, show the opposite pattern. This asymmetry is important. It is less concerning when true targets outrank shuffled decoys, since PLM-based models are expected to assign lower similarity to sequences that violate protein semantics. However, when decoys outrank true targets, homolog detection becomes vulnerable to data-level hijacking in the absence of genuine biological relatedness.

Lastly, the failure mode is illustrated at the level of individual queries and ranking behavior. For non-PLM-based methods, an example query shows that decoy similarities remain close to non-homolog similarities rather than approaching the homolog regime (Figure 5c and Section S2.2.1). In comparison, cumulative target-versus-decoy curves show that PLM-based methods much more readily admit thresholds at which decoys compete with or even outweigh true targets (Figure 5d and Section S2.2.2)). This is precisely the failure mode that the diagnostic is designed to expose, because it indicates that high similarity can be hijacked by synthetic sequence manipulation even in the absence of genuine homology. Taken together, these results show that many PLM-based search methods fail data manipulation safety. Their similarity scores are more susceptible to inflation by shuffled decoys, which means that a high score cannot always be interpreted as reliable evidence of homology when adversarial or composition-matched negatives are present.

### 2.7 PLM-based methods are vulnerable to model-level hijacking by representation perturbation

We finally asked whether high similarity can be induced by manipulating the model itself rather than the input sequences. Following the protocol in Section 4.7, we constructed decoy PLMs by shuffling the weights of the protein language model (*e*.*g*., ESM2) used to produce search embeddings, and then tested whether the resulting representation space could interfere with homolog detection (Figure 5e). This diagnostic applies only to PLM-based methods, because classical alignment-based methods do not rely on learned embedding spaces of this kind. Under this definition, a method satisfies model manipulation safety only if perturbing the representation pipeline does not allow decoy models to compete with or outweigh true homologous matches.

Our analysis shows that current PLM-based search methods are highly vulnerable to this form of manipulation. PLMs with weight perturbation, denoted as decoy PLMs, often rank much more competitively against true targets, as reflected by the deviated AUROC values in Figure 5f. This effect is often more severe than in the data manipulation setting, suggesting that perturbing the PLM can distort homolog detection even more strongly than shuffling the sequence itself. Once the representation space is perturbed, homolog-versus-decoy discrimination becomes substantially less reliable. This behavior indicates that the reported similarity depends strongly on the geometry of the learned embedding space and can be destabilized without introducing any genuine biological signal.

Lastly, the failure mode is illustrated at the level of individual examples. Example queries show that decoy PLMs can assign decoy or non-homolog matches similarities that overlap with, or even exceed, those of true homologs (Figure 5g and Section S2.3). This is a more serious failure than ordinary retrieval noise, because it means that high similarity can be hijacked directly at the model level. Taken together, these results show that many PLM-based search methods fail model manipulation safety, indicating that their similarity scores can be redirected by perturbations of the representation pipeline rather than being stably anchored to biological relatedness.

## 3 Discussion

In this study, we introduced PLM-GUARD, a diagnostic framework for examining what protein search similarity scores mean in the era of language models. Rather than treating search evaluation purely as a ranking problem, we asked whether similarity scores satisfy a set of basic conditions required for biological interpretation. To do so, we organized six controlled sanity checks into three dimensions: biological fidelity, semantic validity, and manipulation safety. Across eight representative search methods, classical alignment-based systems were generally much more stable under these diagnostics, whereas current PLM-based search methods showed broad and systematic failures. These failures were not confined to a single mode of analysis. They appeared in how similarity tracked controlled evolutionary divergence, how it responded to structure-preserving mutation, how it behaved under redundancy and truncation, and how readily it could be inflated by manipulating either the data or the model itself.

This distinction has practical importance. In many downstream settings, sequence search is used not only to retrieve candidates, but also to support annotation transfer, homology claims, and biological reasoning. In such settings, the score itself often carries implicit evidential weight. Our results suggest that this interpretation is much safer for methods whose scores remain anchored to explicit alignment and much less secure for methods whose scores arise primarily from learned embedding geometry. More broadly, PLM-GUARD highlights the need to separate retrieval utility from inferential meaning. A method may still be useful for finding neighbors while producing scores that are unstable under controlled perturbations and therefore difficult to interpret mechanistically. The contribution of PLM-GUARD is not to replace conventional search benchmarks, but to complement them with a diagnostic layer that asks when a similarity score remains biologically meaningful and when that meaning begins to break down.

Last, this study highlights several constructive directions for future extensions. First, a particularly important next step is to develop explicit error control [28] for protein search results, so that returned matches can be accompanied by calibrated confidence rather than raw similarity alone. A natural inspiration comes from target-decoy competition in mass spectrometry proteomics, where decoy matches are used to estimate and control the false discovery rate of reported identifications [29, 30]. An analogous strategy for protein search could use biologically informed decoys or query-specific null constructions to estimate how often high-scoring matches are expected to occur by chance, thereby converting similarity from a heuristic score into an error-controlled discovery signal. Second, future PLM-based search methods could be trained with explicit diagnostic constraints, for example by encouraging monotone behavior under controlled evidence removal, invariance to redundant query content, and robustness to data-level or representation-level perturbations. This would move diagnosis from a purely post hoc exercise toward a constructive design principle. Third, the comparatively good behavior of NEAR suggests that the path forward may lie not in abandoning alignment, but in combining embeddings with alignment-like matching more carefully. Because NEAR still grounds similarity in residue-level correspondence rather than a single global embedding comparison, it preserves more biological grounding than fully PLM-based methods, although its remaining failures also show that hybridization alone is not sufficient. Finally, the proposed sanity checks in this study are intentionally generic. Future work could make them more query aware by stratifying behavior according to sequence length, domain architecture, disorder, taxonomic origin, or low-complexity composition, which may help identify when PLM-based similarity is reliable and when it is especially fragile.

In conclusion, our findings point to a shift in how protein search should be developed and assessed. As learned representations become increasingly central to computational biology, the field will need evaluation frameworks that ask not only whether a method works, but also what its scores mean and under what conditions they can be trusted. We hope that PLM-GUARD provides a useful starting point for that effort. The long-term goal is not merely faster or stronger retrieval, but protein search systems whose similarity scores are interpretable, robust, and reliable enough to support scientific inference.

## 4 Methods

### 4.1 Evaluated protein sequence search methods

BLASTp [1] is the classical protein sequence search method that identifies local alignments through a seed-and-extend strategy and reports alignment-based scores such as bit scores and E-values. It remains a foundational baseline because its scoring framework is closely tied to residue-level substitution models and has long served as a practical standard for sequence similarity search. In our study, BLASTp provides a reference point for methods whose similarity scores have a well-established alignment interpretation. We use BLASTp as an alignment-based comparator and evaluate both its behavior and the diagnostic properties of its reported similarity statistics under our benchmark settings.

MMseqs2 [31] is a fast protein sequence search framework that accelerates large-scale similarity search through efficient indexing, prefiltering, and alignment stages while preserving high sensitivity. It is a key baseline because it is widely used in modern large-database workflows and represents the current high-performance alignment-based paradigm. In our study, MMseqs2 serves as a strong alignment-based reference that emphasizes scalability without abandoning residue-level matching.

DIAMOND [32] is an accelerated protein alignment tool for large-scale sequence search that uses optimized seed-based filtering and extension to achieve high speed while maintaining strong sensitivity. It is a central baseline in our study because it is widely used in modern protein annotation and metagenomic workflows, where throughput and database scale are major constraints. Like BLASTp and MMseqs2, DIAMOND remains alignment-based, which allows direct comparison between accelerated alignment scores and similarity scores from language model based methods.

NEAR [33] is a neural embedding based protein search method designed as a high-speed pre-filter for sensitive homology search. Unlike classical alignment-first methods, NEAR computes learned residue-level embeddings and performs vector search, but unlike general-purpose PLM-based methods, its embeddings are trained specifically to reflect residue relationships supported by trusted sequence alignments. This makes NEAR a relevant intermediate baseline in our study because it combines neural representations with an alignment-oriented training objective and pipeline.

TM-Vec [13] is a PLM-based protein search method that embeds proteins into a structure-aware vector space and retrieves nearest neighbors using vector search. It is trained to predict structural similarity (TM-score) from sequence pairs, which makes it a strong PLM-based baseline for remote homology search and structure-aware search. In the original framework, TM-Vec is used for scalable search, while DeepBLAST is a downstream pairwise alignment module for candidate refinement.

PLMsearch [14] is a PLM-based protein search method for remote homology detection that combines a domain-informed prefilter (PfamClan) with a learned similarity predictor (SS-predictor) trained to approximate structure similarity (TM-score) from protein embeddings. We include PLMSearch because it is a strong and widely cited PLM-based method, which is sensitive protein search beyond classical alignment regimes. It is especially relevant because it combines a practical large-scale pipeline with a score designed to correlate with structure similarity, not just sequence identity.

Deep Homolog Retriever (DHR) [15] is a PLM-based protein search method for protein homology search that uses a dual-encoder architecture and contrastive learning to embed queries and database sequences into a shared vector space. We include DHR because it is a clear example of the dense retrieval paradigm in protein search, where sequence pairs are compared through learned fixed-dimensional embeddings rather than explicit alignment during the search stage. This makes DHR a strong contrast to both classical alignment-first methods and hybrid methods such as NEAR.

DCTdomain [34] is a protein similarity search method that builds domain-aware protein embeddings from PLM residue embeddings by segmenting proteins into domains or subdomains and converting each segment into a compact DCT-based fingerprint. It is an important PLM-based baseline in our study because it explicitly addresses a major weakness of single-vector protein embeddings, namely reduced sensitivity for multidomain proteins and local similarity patterns. In our benchmark, DCTdomain represents a domain-aware embedding retrieval paradigm.

### 4.2 Controlled evolutionary divergence for evolutionary plausibility

Evolutionary plausibility tests whether the similarity scores produced by a sequence search method behave in a biologically coherent way under progressive evolutionary divergence. As mutations accumulate along an evolutionary trajectory, the similarity between a query and its true homolog should decrease. A method that remains faithful to evolutionary relationships should therefore assign systematically lower similarity scores to increasingly diverged variants of the same sequence. Our protocol is designed to test whether this expected monotonic behavior is preserved under controlled sequence evolution.

To generate controlled evolutionary trajectories, we simulated sequence evolution with Pyvolve [35] under the WAG amino acid substitution model [36]. The phylogenetic topology used for simulation is given below and illustrated in Figure 2a:

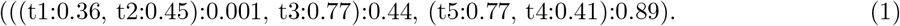

Starting sequences were drawn from the ASTRAL40 database [27], which provides a curated non-redundant collection of protein domains. For each starting sequence, we generated a series of mutated variants corresponding to the nodes of the simulated phylogeny. The original sequence was indexed as mutational stage *k* = 0, and the descendants were indexed by increasing evolutionary divergence. Each variant was then used as a query against the ASTRAL40 database with the evaluated search method. The original sequence at *k* = 0 was retained in the database so that the corresponding true homolog was always present. For each query, we retained the top 1000 retrieval results for analysis.

We then examined how the similarity score assigned to the true homolog changed as mutations accumulated. Let *S*_*k,i*_ denote the similarity score between the query at mutational stage *k* and the homolog corresponding to the *i*-th sequence pair. Under ideal evolutionary behavior, similarity should decrease as divergence increases, so the score at stage *k* should be greater than or equal to the score at stage *k* +1. In practice, however, not all score differences are equally informative. Very small changes may reflect numerical noise or nearly indistinguishable sequences, whereas larger changes provide stronger evidence about whether the similarity function is tracking evolutionary divergence.

To quantify this behavior, we considered stage-to-stage comparisons and ranked them by the absolute magnitude of the score change while retaining the sign that indicates whether the change follows the expected direction. We then evaluated this ranking with the area under the precision-recall curve (AUPRC), treating monotonic decreases as positives and violations as negatives. This metric quantifies how consistently a method assigns larger score changes to comparisons that follow the expected evolutionary trend. Higher AUPRC therefore indicates that the method preserves evolutionary plausibility more faithfully, in the sense that its similarity scores change in a direction and magnitude that better reflect increasing sequence divergence.

### 4.3 Structure-preserving mutation for structural consistency

Structural consistency examines the “twilight zone” of protein homology [23] by testing whether similarity scores remain aligned with structural relatedness when sequence signals are weak. This analysis focuses on remote-homology settings in which sequence identity is low but structural conservation remains substantial. The goal is to determine whether a method is genuinely sensitive to structural relatedness rather than relying on residual sequence similarity or memorized training associations.

To construct a controlled benchmark for this setting, we generated a synthetic dataset of remote homologs with PyRosetta [37]. We began with 1000 template sequences sampled at random from the ASTRAL40 database [27]. For each template, we induced sequence divergence by randomly selecting 60% of residues for mutation. At each selected position, the native amino acid was replaced with a different standard amino acid chosen uniformly at random. After mutation, we applied the Rosetta FastRelax protocol [38] to relax the structure and resolve steric clashes, producing a physically plausible conformation. During mutation and relaxation, side-chain repacking was restricted to mutated residues and their spatial neighbors in order to preserve local topology, and disulfide bonds were explicitly maintained to preserve global structure. Lastly, to ensure that sequence perturbations did not induce large structural deviations, we retained only samples with C*α*-RMSD below 3.0 Å relative to the native structure.

We then validated each mutated structure against its original parent and retained only pairs satisfying standard remote-homology criteria, namely sequence identity below 0.3 and Template Modeling score (TM-score) [39] above 0.5. These filtered pairs formed the synthetic remote-homology set. For each mutated query, we searched against the original ASTRAL40 database and retained all candidates reported by each method. As a natural remote-homology reference, we also queried with the original unmutated sequences and identified ground-truth database hits using the same geometric criteria of sequence identity below 0.3 and TM-score above 0.5.

To quantify structural consistency, we measured the separability between remote homologs as positive samples and non-homologs as negative samples using the area under the receiver operating characteristic curve (AUROC). This metric provides a scale-invariant summary of how well each method preserves structurally meaningful similarity in both the natural and synthetic remote-homology settings. Although AUROC is designed for balanced binary classification, our setting contains many fewer homologs than non-homologs. We therefore repeatedly subsampled the non-homolog set to match the size of the homolog set, computed AUROC on each balanced sample, and reported the mean across 20 repetitions.

### 4.4 Non-informative query augmentation for redundancy stability

Redundancy stability tests whether similarity scores assigned to true homologs remain stable when a query is augmented with redundant or non-informative sequence content. This analysis serves as a control against methods that may inflate similarity because of sequence length or repetitive patterns rather than genuine biological signal. To evaluate this behavior, we generated two augmented variants for each query sequence *q* drawn from the ASTRAL40 database [27]: a “duplicated-self” variant, *q*_dup_ = *q* + *q*, to test sensitivity to redundant information, and a “shuffled-self” variant, *q*_shuf_ = *q* +shuffle(*q*), where shuffle(*q*) denotes a random permutation of *q*, to test sensitivity to appended sequence content that preserves composition but lacks biological meaning.

We then performed similarity searches using both augmented variants and the original sequence as queries against the ASTRAL40 database. For each query, we retained the top 1000 retrieval candidates and extracted the similarity scores assigned to known homologs. These results were aligned into matched pairs (*x*_*i*_, *y*_*i*_), where *x*_*i*_ denotes the score between the original query and a given homolog, and *y*_*i*_ denotes the score between the corresponding augmented query and the same homolog. Pairs were excluded if the homolog did not appear within the top 1000 results for either query version. Under ideal redundancy stability, the augmented query should not systematically alter the homolog score, so that *y*_*i*_ ≈ *x*_*i*_.

To quantify deviations from the expected identity relationship, we defined the Normalized Distance as

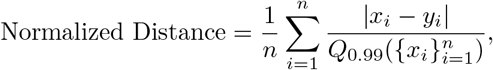

where *n* is the number of retained pairs and 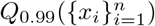 denotes the empirical 99th percentile of the *x*_*i*_ values. This normalization reduces the influence of very small absolute changes, which may reflect numerical noise rather than meaningful instability, and uses the 99th percentile instead of the maximum for greater robustness to outliers. Smaller values of Normalized Distance indicate stronger adherence to the expected invariance under redundant or non-informative query augmentation.

### 4.5 Progressive query truncation for similarity monotonicity

Similarity monotonicity tests whether similarity decreases coherently as informative sequence content is removed from the query. This diagnostic complements redundancy stability by asking not whether scores remain invariant, but whether they attenuate in the expected direction when biological evidence is progressively reduced. To evaluate this behavior, we generated two truncated variants for each query sequence from the ASTRAL40 database by randomly sampling contiguous subsequences that retained exactly 50% and 25% of the original sequence length.

We then searched these truncated variants against the ASTRAL40 database and extracted the similarity scores assigned to true homologs. Using the same matching protocol as in the redundancy stability analysis, we retained homolog pairs for which the same homolog appeared within the top 1000 retrieval candidates for both the longer and shorter query versions. Let *x*_*i*_ denote the score assigned by the longer query and *y*_*i*_ the score assigned by the corresponding shorter query. Under ideal monotonic behavior, removing informative sequence content should not increase similarity, so one expects *y*_*i*_ ≤ *x*_*i*_. To quantify violations of this expected ordering, we defined the Monotonicity Loss as:

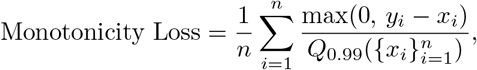

where *n* is the number of retained homolog pairs and 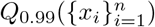 denotes the empirical 99th percentile of the scores from the longer query. Because we used two truncated variants that retained exactly 50% and 25% of the original sequence length, we reported the monotonicity loss as the sum across these two settings.

This loss is zero when similarity decreases or remains unchanged after truncation, and becomes positive only when similarity increases despite the removal of informative sequence content. The normalization reduces the influence of numerical noise from very small score differences and improves robustness by using the 99th percentile, which can be less sensitive to outliers. Larger values of Monotonicity Loss therefore indicate stronger violations of similarity monotonicity.

### 4.6 Shuffled decoys for data manipulation safety

Data manipulation safety tests whether a similarity search method assigns spuriously elevated scores to synthetic sequences that preserve low-level statistical properties while lacking genuine homology. To assess data-level vulnerability, we constructed a decoy counterpart for each sequence in the ASTRAL40 dataset by randomly shuffling its amino acid residues. This operation destroys local sequence motifs and structural coherence while preserving global amino acid composition and sequence length. Pooling these shuffled decoys with the original sequences produced a combined target database that challenges each method to distinguish true biological signal from statistically matched but biologically meaningless noise.

We then searched each original ASTRAL40 sequence against this combined target database and retained all returned hits. Retrieved sequences were partitioned into three categories based on the homology criteria used throughout the study. True homologs were defined by shared SCOP family or superfamily labels. All other hits were treated as negatives and further separated into natural negatives, consisting of non-homologous original sequences, and synthetic negatives, consisting of shuffled decoy sequences.

To summarize score behavior, we first visualized the global separation among these groups with kernel density estimation, comparing the score distributions of true homologs, natural negatives, and shuffled decoys. We then quantified target-versus-decoy discrimination with AUROC, computed separately for each query. In this analysis, both true homologs and non-homologs were treated as the positive class, while shuffled decoys were treated as the negative class. Under this setup, AUROC reflects whether biologically meaningless decoy sequences are spuriously assigned higher scores than genuine targets, rather than measuring general retrieval quality alone. Comparing AUROC against natural negatives and against shuffled decoys therefore reveals whether a method is specifically vulnerable to data-level manipulation by statistically matched but non-homologous sequences.

### 4.7 Perturbed PLMs for model manipulation safety

Model manipulation safety tests whether similarity scores remain stable when the learned representation pipeline is perturbed in a controlled way. Unlike data manipulation safety, which alters the input sequences, this analysis asks whether homolog detection can be distorted by manipulating the model while leaving the biological sequences unchanged. A method satisfies model manipulation safety only if such perturbations do not allow decoy representations to compete with or outweigh genuine homologous matches. This diagnostic is specific to PLM-based search methods, because classical alignment-based methods do not rely on learned embedding spaces that can be perturbed in this manner. It therefore isolates a failure mode unique to embedding-based protein search, namely that high similarity may arise from instability in learned geometry rather than from genuine biological relatedness.

To evaluate the vulnerability to model-level manipulation, we constructed a decoy version of each applicable PLM by permuting the final fully connected layer used to produce search embeddings. For PLMSearch, we permuted the final fully connected layer of ESM-1b model by shuffling the columns of its weight matrix. For TM-Vec, we applied the same perturbation to the final fully connected layer of the ProtTrans model. For DHR, we permuted the final fully connected layer of ESM-2b model in the same way. DCTdomain differs from these methods because it extracts protein language model representations from two internal layers of ESM2-t33-650M, specifically layers 14 and 20. Accordingly, we permuted the fully connected layer in both of these layers. In each case, the perturbation preserves the overall weight distribution while disrupting the learned correspondence between internal representations and the final embedding space used for sequence search.

Using these decoy PLMs, we repeated the search procedure against the same ASTRAL40 database and compared the resulting scores with those obtained from the original PLMs. As in the data manipulation safety analysis, homologous targets were treated as the positive class and decoy or non-homolog matches were treated as the negative class. We then quantified target-versus-decoy discrimination with AUROC, computed separately for each query. Under this setup, AUROC has the same interpretation as in the data manipulation setting. It reflects the extent to which the relative ranking between targets and negatives is altered under PLM perturbation. More deviated AUROC indicates that similarity has become more vulnerable to model-level hijacking and less reliable as biological evidence.

## 5 Data Availability

All datasets used for evaluating the models are available via Zenodo at https://doi.org/10.5281/zenodo.19795993.

## 6 Code Availability

The code used to reproduce the analyses, figures, and evaluation protocols in this study is publicly available at https://github.com/batmen-lab/PLMGuard. The repository includes implementations of the six sanity checks in PLM-GUARD, scripts for running the evaluated search methods, and code for reproducing the reported results.

## 7 Acknowledgements

This research was supported by startup funding from the University of Wisconsin–Madison RISE AI Initiative.

## 8 Author contributions

H.Z. implemented the code, set up and preprocessed the datasets, and performed the analysis.. Y.Y. participated the analysis. H.Z. and Y.Y.L. prepared the figures and participated in the discussion. Y.Y.L. wrote the manuscript. All authors participated the discussion. All authors reviewed the manuscript.

## References

[1] Altschul, S. F., Gish, W., Miller, W., Myers, E. W. & Lipman, D. J. A basic local alignment search tool. Journal of Molecular Biology 115, 403–410 (1990).

[2] Altschul, S. F. et al. Gapped BLAST and PSI-BLAST: A new generation of protein database search programs. Nucleic Acids Research 15, 3389–3402 (1997).

[3] Pearson, W. R. & Lipman, D. J. Improved tools for biological sequence comparison. Proceedings of the National Academy of Sciences 85, 2444–2448 (1988).

[4] Lu, Y. Y., Noble, W. S. & Keich, U. A BLAST from the past: revisiting blastp’s E-value. Bioinformatics btae729 (2024).

[5] Smith, T. & Waterman, M. Identification of common molecular subsequences. Journal of Molecular Biology 147, 195–197 (1981).

[6] Eddy, S. R. Accelerated profile HMM searches. PLoS Computational Biology 7, e1002195 (2011).

[7] Pearson, W. R. An introduction to sequence similarity (“homology”) searching. Current Protocols in Bioinformatics 41, 3–1 (2013).

[8] Durbin, R., Eddy, S., Krogh, A. & Mitchison, G. Biological Sequence Analysis (Cambridge UP, 1998).

[9] Jr, R. L. D. Sequence comparison and protein structure prediction. Current Opinion in Structural Biology 1?, 374–384 (2006).

[10] Rives, A. et al. Biological structure and function emerge from scaling unsupervised learning to 250 million protein sequences. Proceedings of the National Academy of Sciences of the United States of America 118, e2016239118 (2021).

[11] Xiao, Y. et al. Protein large language models: A comprehensive survey. arXiv preprint 2502.17504 (2025).

[12] Lin, Z. et al. Evolutionary-scale prediction of atomic-level protein structure with a language model. Science 379, 1123–1130 (2023).

[13] Hamamsy, T. et al. Protein remote homology detection and structural alignment using deep learning. Nature Biotechnology 41, 975–985 (2024).

[14] Liu, W. et al. PLMSearch: Protein language model powers accurate and fast sequence search for remote homology. Nature Communications 15, 2775 (2024).

[15] Hong, L. et al. Fast, sensitive detection of protein homologs using deep dense retrieval. Nature Biotechnology 1–13 (2024).

[16] Bileschi, M. L. et al. Using deep learning to annotate the protein universe. Nature Biotechnology 40, 932–937 (2022).

[17] Heinzinger, M. et al. Modeling aspects of the language of life through transfer-learning protein sequences. BMC Bioinformatics 10, 723 (2019).

[18] Vig, J. et al. BERTology meets biology: Interpreting attention in protein language models. International Conference on Learning Representations (2021).

[19] Unsal, S. et al. Learning functional properties of proteins with language models. Nature Machine Intelligence 4, 227–245 (2022).

[20] Liu, Y. et al. Benchmarking protein sequence and structure search methods for remote homology detection (2026).

[21] Doshi-Velez, F. & Kim, B. Towards a rigorous science of interpretable machine learning. arXiv preprint 1702.08608 (2017).

[22] Chothia, C. & Lesk, A. M. The relation between the divergence of sequence and structure in proteins. EMBO Journal 5, 823–826 (1986).

[23] Rost, B. Twilight zone of protein sequence alignments. Protein Engineering 11, 85–94 (1999).

[24] Karlin, S. & Altschul, S. F. Methods for assessing the statistical significance of molecular sequence features by using general scoring schemes. Proceedings of the National Academy of Sciences of the United States of America 87, 2264–2268 (1990).

[25] Elias, J. E. & Gygi, S. P. Target-decoy search strategy for mass spectrometry-based proteomics. Methods in Molecular Biology 04(2010).

[26] Goodfellow, I. J., Shlens, J. & Szegedy, C. Explaining and harnessing adversarial examples. International Conference on Learning Representations (2014).

[27] Fox, N. K., Brenner, S. E. & Chandonia, J.-M. SCOPe: Structural classification of proteins—extended, integrating SCOP and ASTRAL data and classification of new structures. Nucleic Acids Research 41, D304–D309 (2014).

[28] Benjamini, Y. & Yekutieli, D. The control of the false discovery rate in multiple testing under dependency. The Annals of Statistics 19, 1165–1188 (2001).

[29] Elias, J. E. & Gygi, S. P. Target-decoy search strategy for increased confidence in large-scale protein identifications by mass spectrometry. Nature Methods 4, 207–214 (2007).

[30] Gupta, N., Bandeira, N., Keich, U. & Pevzner, P. Target-decoy approach and false discovery rate: When things may go wrong. Journal of the American Society for Mass Spectrometry 11, 1111–1120 (2011).

[31] Steinegger, M. & Söding, J. Mmseqs2 enables sensitive protein sequence searching for the analysis of massive data sets. Nature Biotechnology 35, 1026–1028 (2017).

[32] Buchfink, B., Reuter, K. & Drost, H. G. Sensitive protein alignments at tree-of-life scale using DIAMOND. Nature Methods 18, 366–368 (2021).

[33] Olson, D. et al. NEAR: neural embeddings for amino acid relationships. Bioinformatics 41, i449–i457 (2025).

[34] Iovino, B. G., Tang, H. & Ye, Y. Protein domain embeddings for fast and accurate similarity search. Genome Research 34, 1434–1444 (2024).

[35] Spielman, S. J. & Wilke, C. O. Pyvolve: a flexible Python module for simulating sequences along phylogenies. PloS One 10, e0139047 (2015).

[36] Whelan, S. & Goldman, N. A general empirical model of protein evolution derived from multiple protein families using a maximum-likelihood approach. Molecular Biology and Evolution 18, 691–699 (2001).

[37] Chaudhury, S., Lyskov, S. & Gray, J. Pyrosetta: a script-based interface for implementing molecular modeling algorithms using Rosetta. Bioinformatics 1?, 689–691 (2010).

[38] Tyka, M. D. et al. Alternate states of proteins revealed by detailed energy landscape mapping. Journal of Molecular Biology 405, 607–618 (2011).

[39] Zhang, Y. & Skolnick, J. Scoring function for automated assessment of protein structure template quality. Proteins 57, 702–710 (2004).

