## Supplement for "Diagnosing protein sequence search in the era of language models"

### S1 Protein sequence search methods

#### S1.1 BLASTp (Protein BLAST)

BLASTp [Altschul et al., 1990] is a widely used local protein sequence alignment method and serves as a canonical alignment-based baseline in this study. We include BLASTp because it represents the historical and methodological reference for sequence similarity search, with scores derived from residue-level alignments and statistical significance estimates that are routinely used in biological inference. This makes BLASTp particularly important that evaluates not only protein search quality but also the interpretability and reliability of similarity scores.

BLASTp uses a heuristic seed-and-extend procedure for local alignment. It first identifies short high-scoring word matches between a query and target sequence, then extends these seeds to form longer local alignments, and finally computes alignment scores under a substitution matrix and gap penalty scheme. BLASTp reports several outputs, including raw alignment score, bit score, and E-value. In practice, the E-value is often interpreted as a significance-like measure of how likely a match of similar quality would occur by chance in the searched database, while the bit score provides a normalized alignment strength. We ran BLASTp using the NCBI BLAST+ implementation [Camacho et al., 2009] with a fixed set of search parameters across all benchmark tasks. The substitution matrix, gap opening and extension penalties, compositional adjustment settings, low-complexity masking options, and hit reporting thresholds were kept constant unless otherwise noted. We used a common target database for all methods and retained BLASTp outputs needed for both protein search evaluation and score diagnostics.

For ranking-based evaluation, we used the native BLASTp ranking induced by alignment score statistics. For diagnostic analyses focused on score interpretability, we tracked the method's reported similarity statistics, including bit score and E-value, and examined how these quantities behave under the controlled perturbations and benchmark conditions defined in our study. When multiple alignments were reported for the same query target pair, we used the top-scoring alignment. We did not reinterpret BLASTp scores beyond their native definitions. Instead, we used them directly and assessed their behavior under a unified diagnostic protocol, allowing a method-agnostic comparison of score interpretability across alignment-based and language model based search paradigms.

#### S1.2 MMseqs2

MMseqs2 [Steinegger and Söding, 2017] is a high-throughput sequence search and clustering framework designed for large-scale protein analysis. We include MMseqs2 as a baseline because it is one of the most widely used modern alignment-based tools for sensitive and scalable protein search. It is especially relevant to this study because it occupies an important middle ground between classical alignment methods and newer embedding-based search systems, combining alignment-based interpretability with aggressive engineering for speed.

MMseqs2 uses a multistage search pipeline that typically includes a fast prefilter to identify promising candidates followed by more detailed alignment evaluation. The prefilter stage reduces the search space substantially, and subsequent stages refine candidate matches using alignment-based scoring. This design enables MMseqs2 to achieve high throughput while maintaining strong sensitivity in many practical settings. The method reports alignment-related similarity measures, and the final ranking is based on native scoring statistics produced by the search pipeline. We ran MMseqs2 with a single standardized configuration for all benchmark tasks unless a task required a predefined variant. Key settings include the search mode, sensitivity parameter, alignment mode, E-value threshold, maximum reported hits, and thread count. We preserved the same target database and hit filtering rules across methods. When MMseqs2 offered optional accelerated or approximate settings, we documented whether these were enabled, as such choices can affect both runtime and score behavior.

We used MMseqs2's native ranking outputs for protein search evaluation and retained the reported similarity statistics for diagnostic analysis. In particular, we examined how the method's alignment-derived scores behave across our diagnostic tasks rather than treating the protein search rank as the only outcome. When multiple hits were reported for the same pair under different alignment variants, we retained the top result according to the native MMseqs2 ranking criterion. The goal in our study is not to force score equivalence across methods, but to compare whether each method's native similarity statistics remain interpretable and stable under the same controlled diagnostic conditions.

#### S1.3 DIAMOND

DIAMOND [Buchfink et al., 2021] is a fast protein sequence aligner developed for high-throughput sequence search, especially in settings with very large reference databases. We include DIAMOND because it is one of the most influential accelerated alignment-based methods in contemporary computational biology and is frequently used as a practical substitute for BLAST-like protein search in large-scale applications. For a diagnosis focused on the meaning and reliability of similarity scores, DIAMOND is essential because it preserves alignment-based scoring while introducing substantial algorithmic acceleration.

DIAMOND follows the general alignment-based search paradigm but incorporates optimized indexing, seed processing, and extension heuristics to improve speed and scalability. It uses a seed-and-extend workflow with high-performance filtering and alignment refinement, enabling rapid candidate generation and scoring against large protein databases. The method reports alignment-related scores and significance statistics, which makes it suitable for direct comparison with other alignment-based baselines and for diagnostic analyses of score behavior. We ran DIAMOND with a standardized configuration across benchmark tasks, including fixed sensitivity mode, score thresholds, maximum target reporting settings, and multithreading parameters. Because DIAMOND offers multiple sensitivity presets and search tradeoffs, we explicitly report the chosen mode and keep it constant in all experiments unless stated otherwise. We retained native output fields required for both ranking evaluation and score diagnostics.

We examined the method's reported alignment-based similarity statistics, including significance-related outputs where available, to assess how they behave under the same perturbation and control regimes used for other methods. When DIAMOND returned multiple high-scoring segment pairs for a query target pair, we used the top-scoring result under the native ranking criterion. We analyzed DIAMOND in its native form and compared its score behavior to other baselines under a shared diagnostic framework.

#### S1.4 NEAR

NEAR (Neural Embeddings for Amino acid Relationships) [Olson et al., 2025] is a neural representation learning method for protein search introduced as a fast and sensitive prefilter for homology detection. We include NEAR as a baseline because it occupies an important methodological bridge between classical alignment-based search and PLM-based search. NEAR uses vector embeddings and nearest-neighbor search, which places it in the embedding-based search family, but it is trained with guidance from trusted sequence alignments and is designed to recover alignment-relevant candidates rather than general sequence semantics. This makes NEAR particularly informative for our study, by disentangling protein search performance from the statistical meaning and reliability of similarity scores.

NEAR computes residue-level embeddings for both query and target proteins and stores target residue embeddings in a vector index. Given a query, it performs residue-level nearest-neighbor search and then aggregates neighbor evidence to score candidate target sequences. The model is trained with contrastive learning using trusted sequence alignments so that residues likely to align to one another are placed close in embedding space. This differs from classical alignment tools, which score sequence similarity directly through substitution matrices and gap models, and also differs from many PLM-based search methods, which often use generic sequence embeddings not trained specifically for homology filtering. In this sense, NEAR is neural in representation but alignment-oriented in supervision and downstream intent. We ran NEAR using its default search pipeline unless otherwise noted. NEAR uses a compact ResNet-based embedding model and a FAISS vector search backend for efficient nearest-neighbor search over residue embeddings. Because NEAR is intended as a pre-filter and not a full alignment engine, we used its native search outputs directly for ranking and diagnostic evaluation rather than inserting an external alignment reranking stage. Any changes to embedding dimensionality, index type, quantization, or search parameters were fixed across experiments and reported explicitly to ensure reproducibility.

We used NEAR's native sequence-level search score, which is produced by aggregating residue-level nearest-neighbor similarity evidence, for ranking and downstream diagnostic analysis. Importantly, this score is not an E-value and is not calibrated as a classical alignment significance statistic. We therefore treat it as a method-native similarity score and evaluate its ranking utility and diagnostic behavior separately from statistical significance metrics used in alignment-based tools. This distinction is important because one of our goals is to compare protein search quality and score interpretability as separate axes. We used NEAR in its native form and did not post hoc calibrate its scores to alignment-based statistics. Our comparison therefore focuses on a method-agnostic diagnostic question: how well each method's native similarity outputs support reliable ranking and interpretation under a shared benchmark protocol.

#### S1.5 TM-Vec

TM-Vec [Hamamsy et al., 2024] is a structure-aware protein search method introduced as the search component of a two-stage pipeline, with DeepBLAST providing pairwise alignments after the search. We include TM-Vec as a core PLM-based baseline because it is explicitly designed for scalable remote homology search using sequence-only input while optimizing for structural similarity rather than sequence identity. This is directly relevant to our paper because it represents a modern PLM-based search paradigm with a learned similarity objective that differs fundamentally from classical alignment scores.

TM-Vec uses a pretrained PLM (ProtTrans/ProtT5 in the original work) to generate residue embeddings, then applies a learned twin-network encoder to map each protein to a fixed-length vector. The model is trained on pairs of proteins with known structures so that the cosine similarity between protein vectors approximates the TM-score, a structural similarity metric. At inference time, target proteins are embedded offline and indexed, and queries are retrieved by nearest-neighbor search in embedding space. In the original paper, this stage is followed by DeepBLAST to produce pairwise structural alignments for top candidates, but TM-Vec itself is the search engine.

We used TM-Vec's native similarity output, which is a learned predictor of structural similarity (TM-score-like behavior) derived from embedding-space similarity. This score is not a classical alignment significance statistic such as an E-value. We therefore treat it as a method-native protein search score and analyze its ranking behavior and score stability separately from alignment-based tools. If DeepBLAST refinement was applied in selected analyses, its alignment outputs were reported as a separate stage. Our comparison focuses on whether each method's native similarity outputs support reliable ranking and interpretation under a shared diagnostic protocol.

#### S1.6 PLMsearch

PLMsearch [Liu et al., 2024] is a sequence-only homolog search method that uses deep sequence representations from a pretrained PLM and a learned structural-similarity predictor to improve remote homology detection. We include PLMSearch because it is a strong and widely cited PLM-based search baseline, which is sensitive protein search beyond classical alignment regimes. It is especially relevant because it combines a practical large-scale protein search pipeline with a score designed to correlate with structure similarity, not just sequence identity.

PLMSearch consists of three stages. First, PfamClan uses Pfam clan annotations to prefilter candidate targets for each query. Second, a PLM generates embeddings, and SS-predictor predicts a similarity score for each candidate pair using a learned predictor trained against structure similarity. Third, PLMSearch ranks candidate pairs by the predicted similarity score and outputs results per query. PLMAlign is also described, which is a separate alignment module that uses per-residue embeddings to build an embedding-derived substitution matrix for Smith–Waterman or Needleman–Wunsch alignment, but PLMSearch itself is the protein search component.

We used the native PLMSearch ranking score produced by SS-predictor. This score is a learned similarity quantity intended to reflect structural similarity and is not a classical alignment significance statistic such as an E-value. We therefore treat it as a method-native score and analyze its ranking behavior and diagnostic properties separately from alignment-based statistics. Our goal is to compare how well each method's own similarity outputs support reliable ranking and interpretation under a shared diagnostic protocol.

#### S1.7 Deep Homolog Retriever

Deep Homolog Retriever (DHR) [Hong et al., 2024] is an ultrafast homolog detection method that combines a protein language model with dense retrieval techniques. We include DHR because it is a clear example of the dense retrieval paradigm in protein search, where sequence pairs are compared through learned fixed-dimensional embeddings rather than explicit alignment during the protein search stage. This makes DHR a strong contrast to both classical alignment-first methods and hybrid methods such as NEAR.

DHR uses a dual-encoder (bi-encoder) architecture with separate query and candidate encoders, both initialized from a PLM. The model is trained with contrastive learning so that homologous pairs are mapped close together while non-homologous pairs are pushed apart in embedding space. At inference time, database sequences are embedded offline, query embeddings are computed online, and similarity is computed via dot product, which supports efficient top-k search with approximate nearest-neighbor indexing (FAISS). The original paper also describes a teacher-student style construction where homologous pairs are derived from MSA-based pipelines (for example JackHMMER/HHblits-derived supervision) during training.

We used DHR's native embedding-space similarity score (dot product between query and candidate embeddings). This score is not an E-value and is not directly tied to explicit residue-level alignment. We treat it as a method-native protein search score and analyze its ranking and diagnostic behavior separately from alignment-derived statistics. This lets us compare score reliability and interpretability across methods without imposing artificial score harmonization.

#### S1.8 DCTdomain

DCTdomain [Iovino et al., 2024] is a PLM-based protein similarity method that augments whole-protein embeddings with domain-level embeddings. We include DCTdomain because it addresses an important practical issue in embedding-based search, where per-protein pooled embeddings can miss local or domain-specific homology, especially in multidomain proteins. The method is directly relevant to our diagnostic framework because it changes not only the embedding representation but also the granularity of similarity scoring.

DCTdomain uses residue embeddings and contact map predictions from ESM-2 to segment proteins into domains or subdomains. It then applies a discrete cosine transform (DCT) to residue embeddings within each whole protein and each predicted domain to produce compact DCT fingerprints. The method reports two scores, a whole-protein score (DCTglobal) and a domain-aware score (DCTdomain) that uses all fingerprints from the full protein and predicted domains. Domain segmentation is performed using a recursive algorithm (RecCut) designed to reduce runtime from cubic to quadratic complexity with respect to protein length.

We fixed the domain segmentation and fingerprinting settings across experiments. We retained the method's native domain-aware similarity scoring and did not replace the domain segmentation module with external annotations. If both DCTglobal and DCTdomain scores were available, we report which score was used in each analysis and keep this choice fixed within each benchmark. This preserves the method's intended behavior and allows a fair comparison of score interpretability across different PLM-based methods.

### S2 Evaluation of evolutionary plausibility

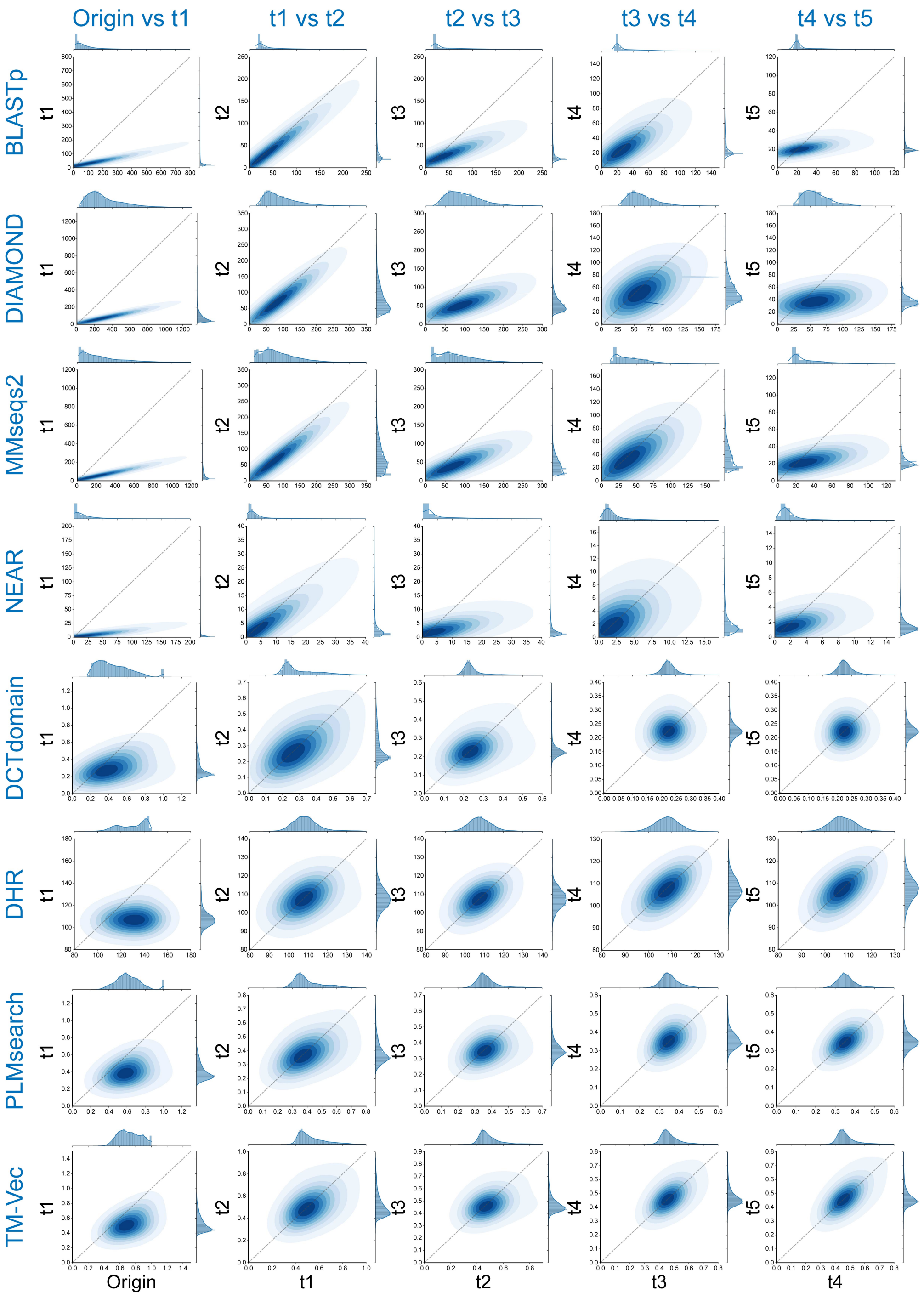

Figure S1: **Evaluation of evolutionary plausibility across protein search methods.**

##### S2.1 Evaluation of structural consistency

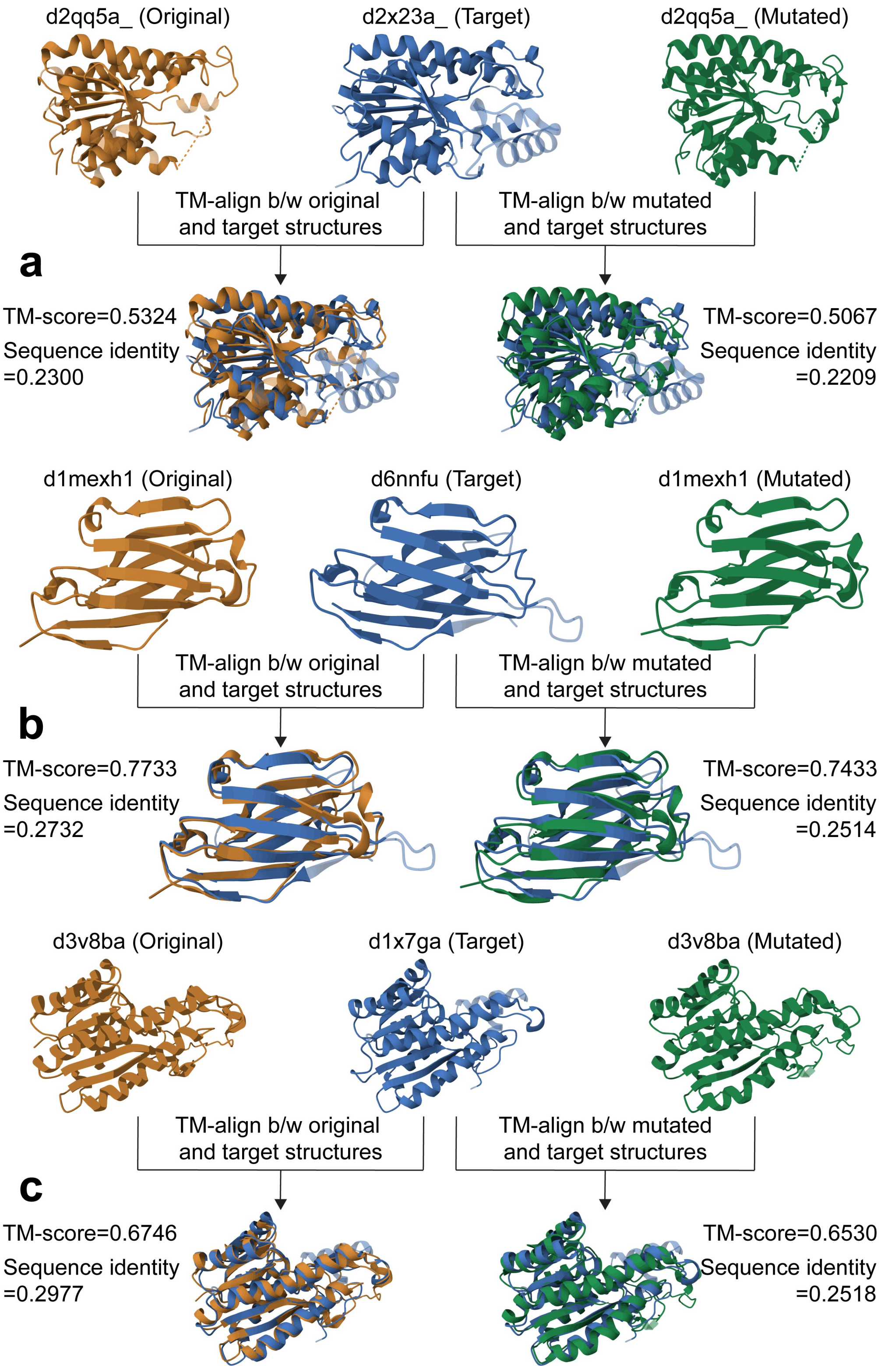

Figure S2: **More examples of remote homolog pairs for structural consistency evaluation.**

#### S2.2 Evaluation of data manipulation safety

##### S2.2.1 Comparison of non-homolog and decoy similarity distributions

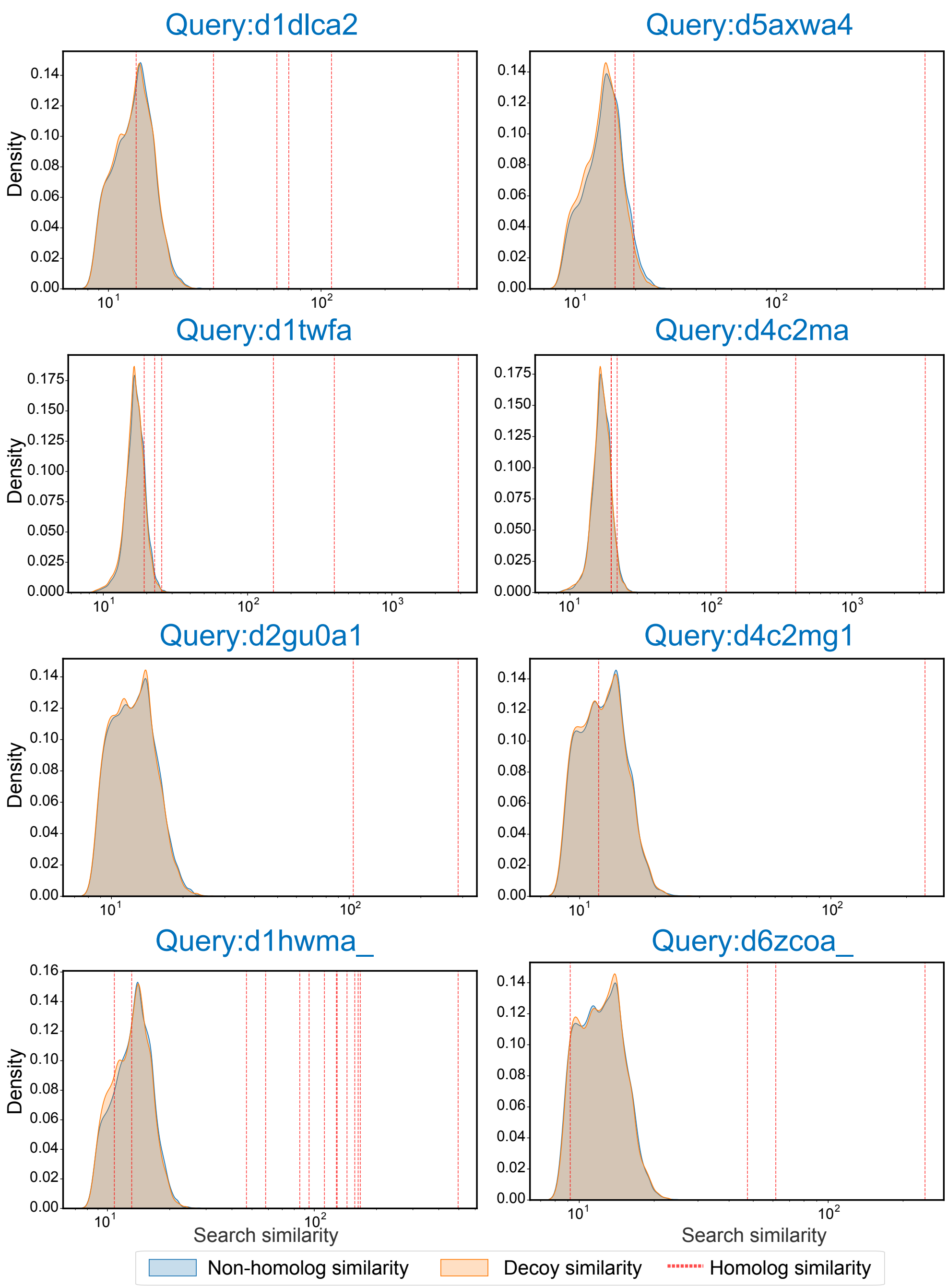

Figure S3: **Comparison of non-homolog and decoy similarity distributions across representative queries using BLASTp.**

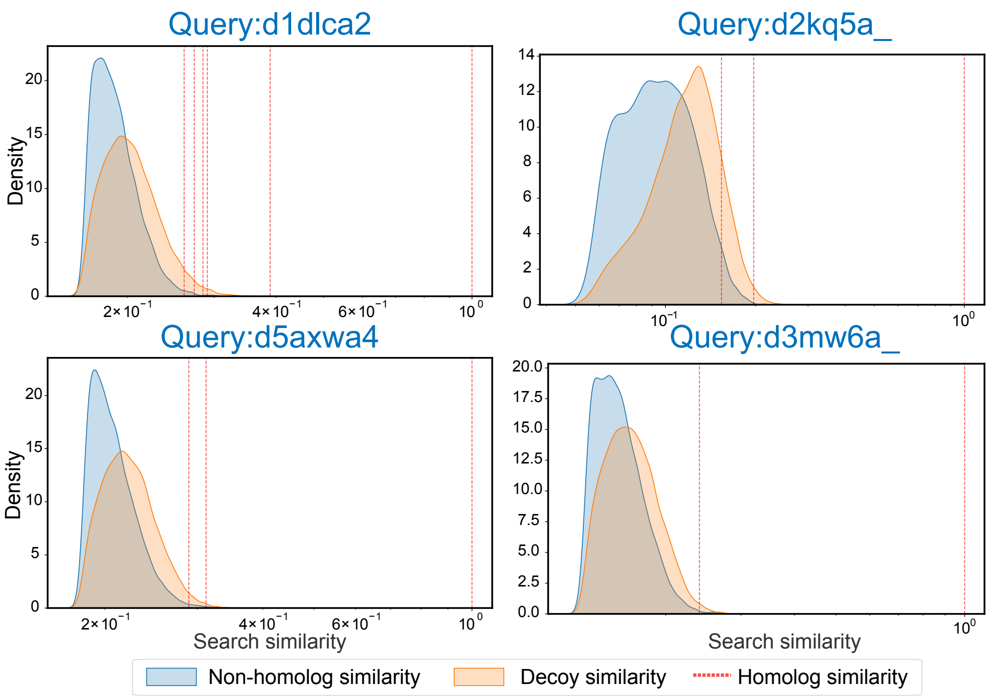

Figure S4: **Comparison of non-homolog and decoy similarity distributions across representative queries using DCTdomain.**

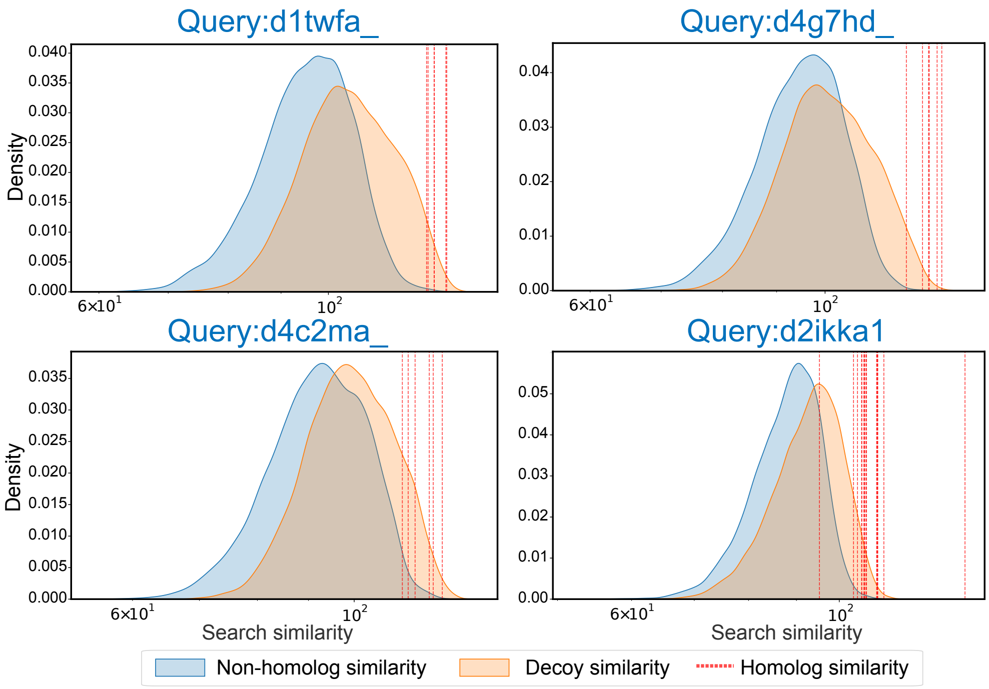

Figure S5: **Comparison of non-homolog and decoy similarity distributions across representative queries using DHR.**

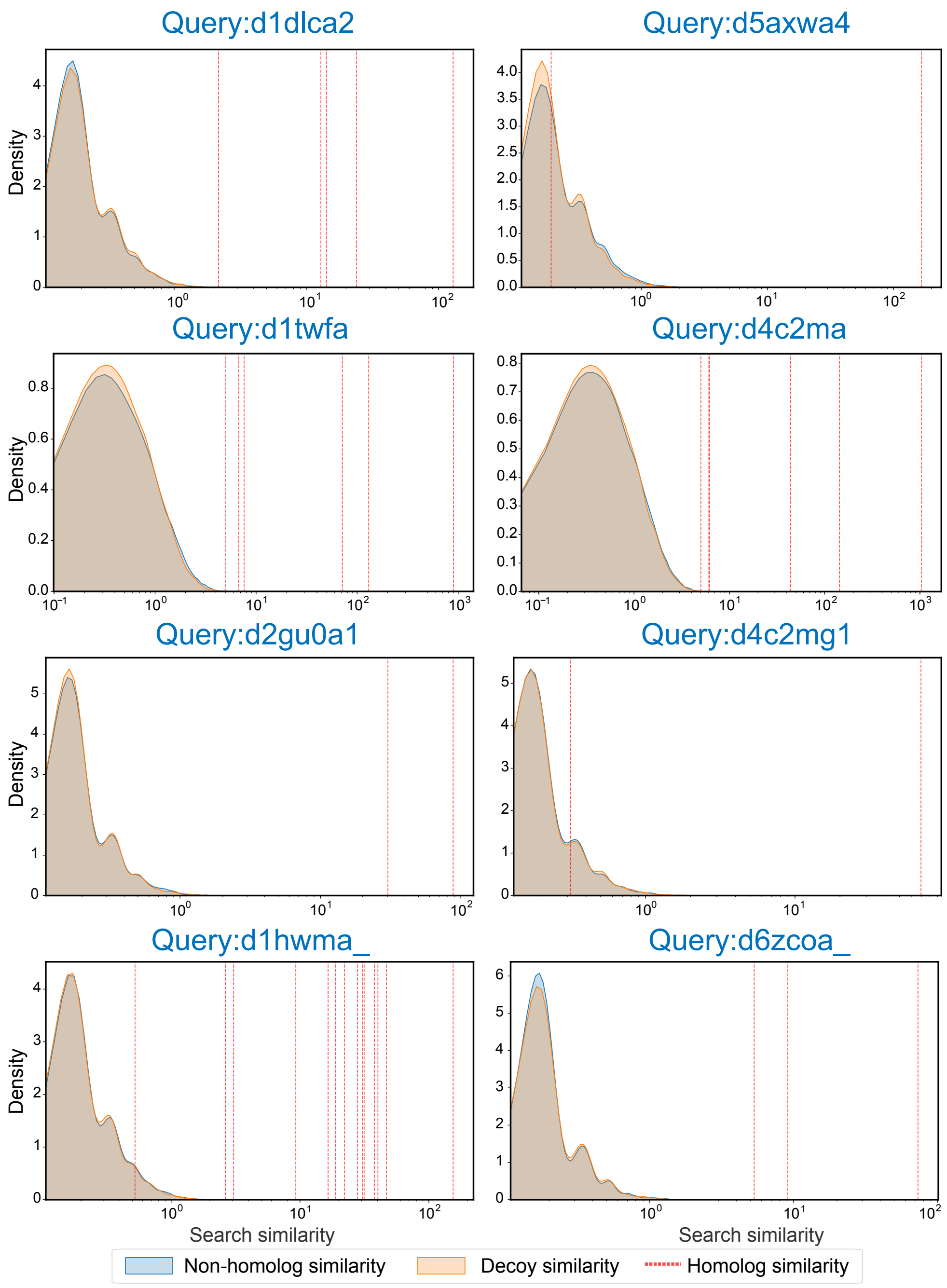

Figure S6: **Comparison of non-homolog and decoy similarity distributions across representative queries using NEAR.**

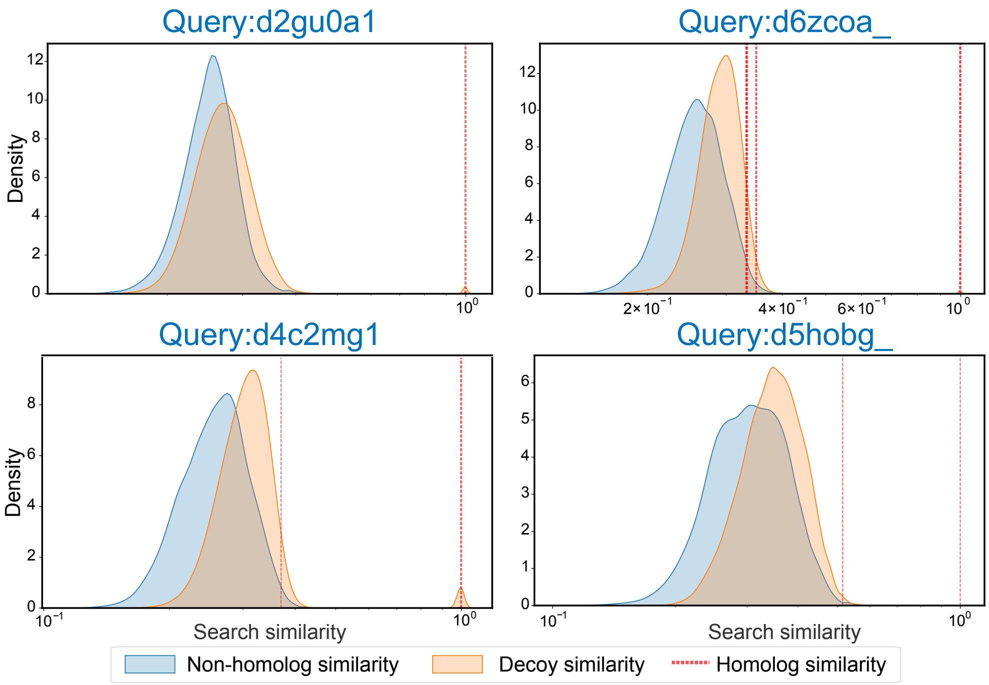

Figure S7: **Comparison of non-homolog and decoy similarity distributions across representative queries using PLMSearch,.**

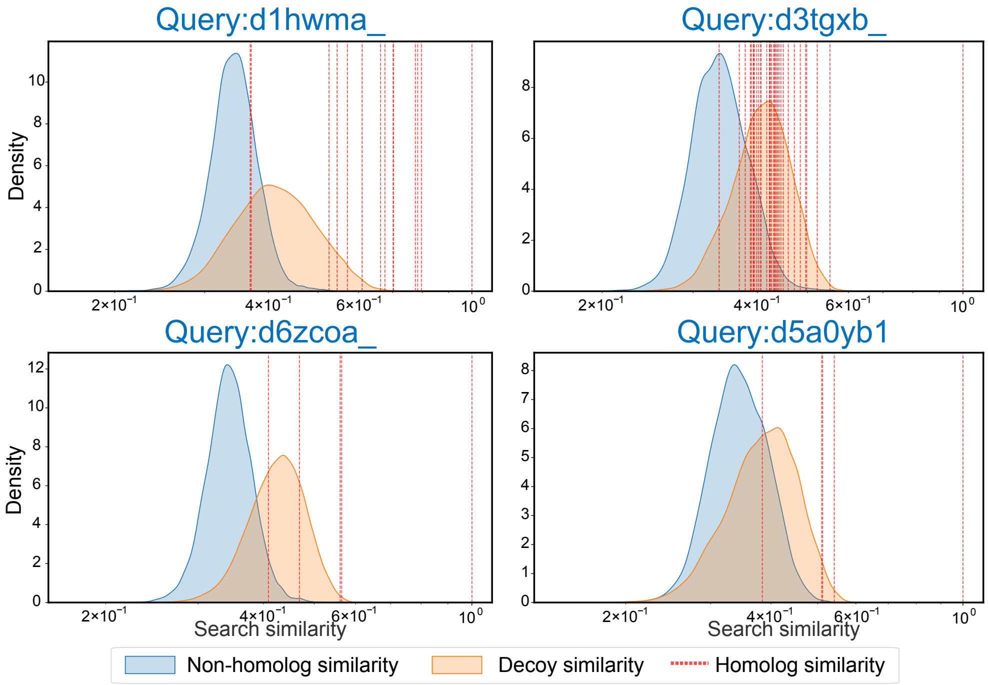

Figure S8: **Comparison of non-homolog and decoy similarity distributions across representative queries using TM-Vec.**

##### S2.2.2 Comparison of cumulative target and decoy fractions

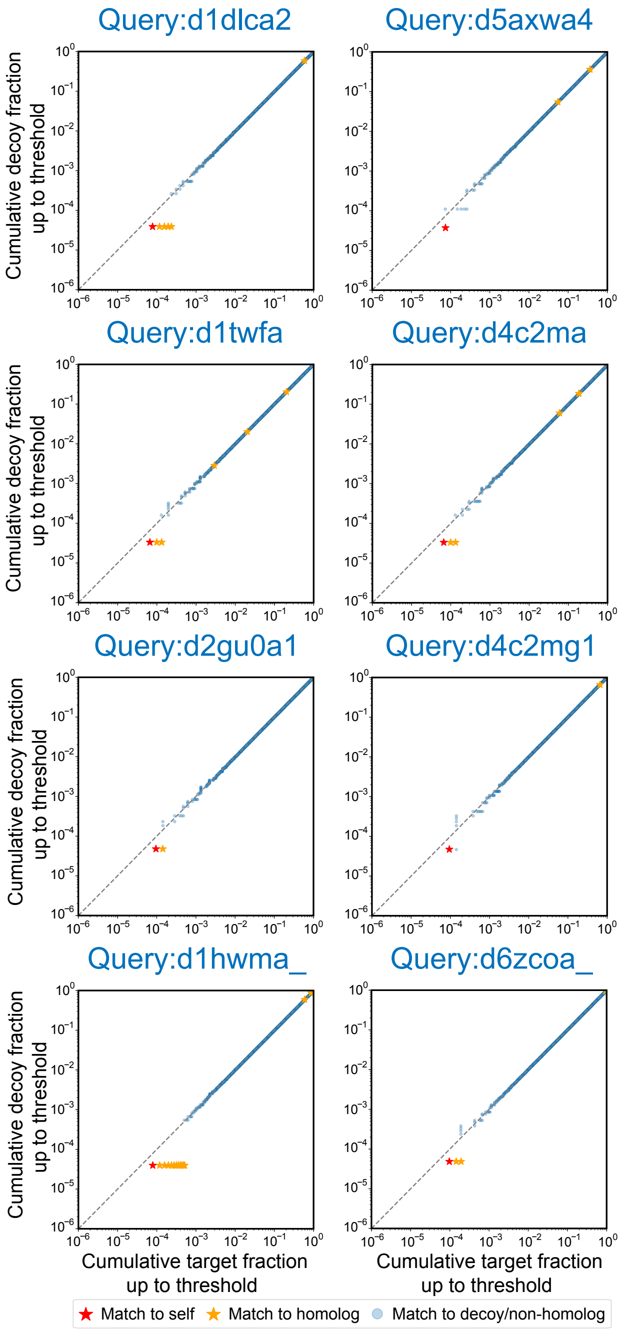

Figure S9: **Comparison of cumulative target and decoy fractions across representative queries using BLASTp.**

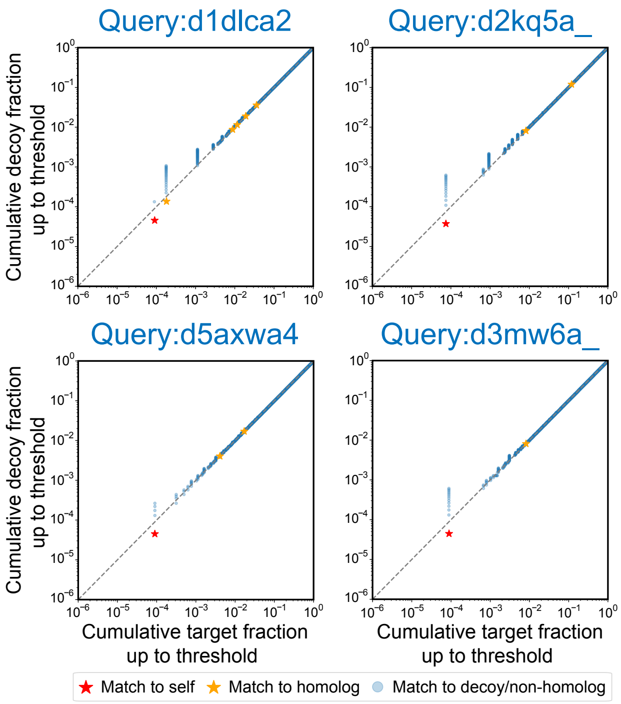

Figure S10: **Comparison of cumulative target and decoy fractions across representative queries using DCTdomain.**

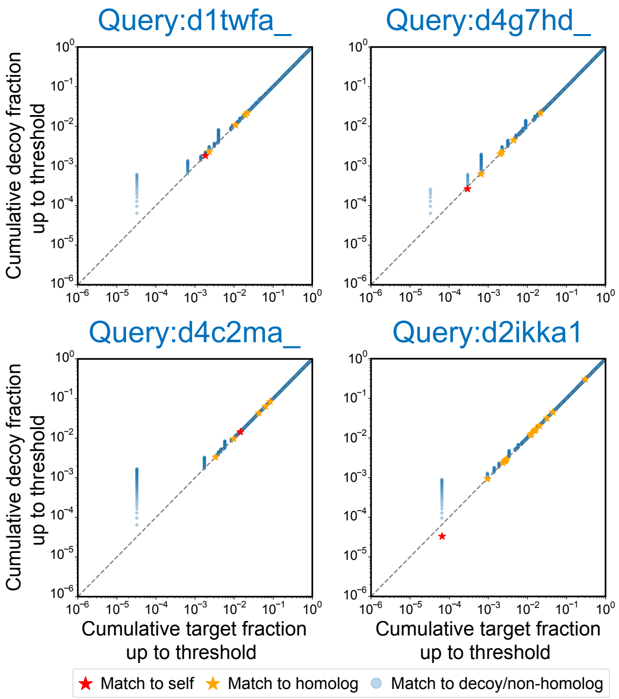

Figure S11: **Comparison of cumulative target and decoy fractions across representative queries using DHR.**

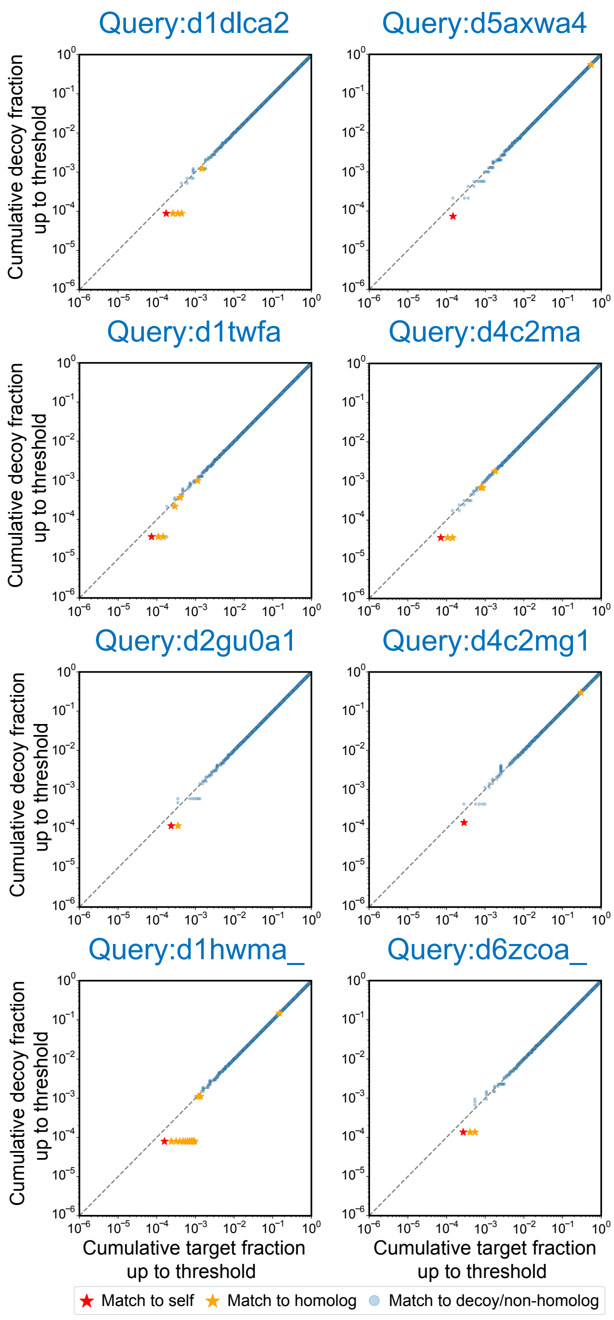

Figure S12: **Comparison of cumulative target and decoy fractions across representative queries using NEAR.**

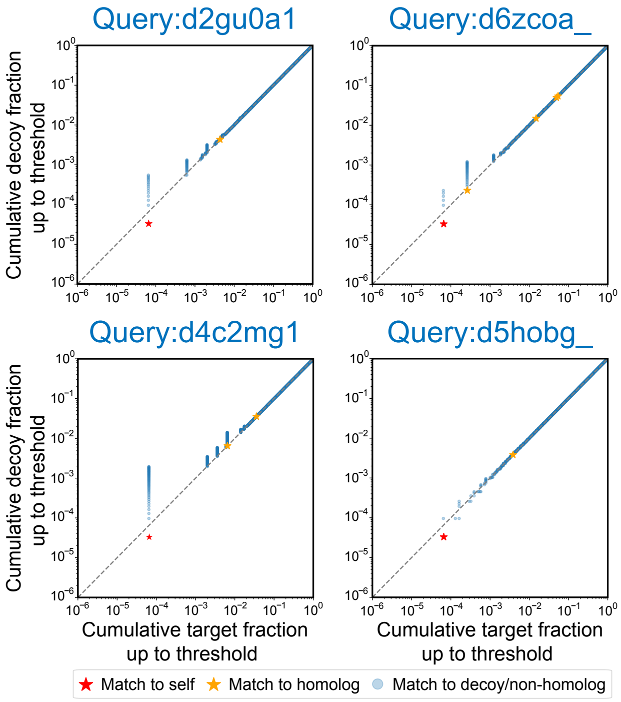

Figure S13: **Comparison of cumulative target and decoy fractions across representative queries using PLMSearch.**

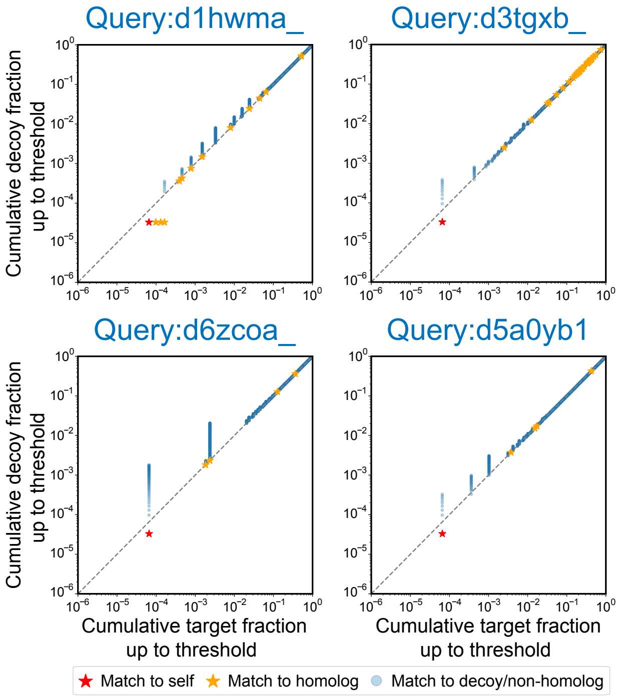

Figure S14: **Comparison of cumulative target and decoy fractions across representative queries using TM-Vec.**

#### S2.3 Evaluation of model manipulation safety

##### S2.3.1 Comparison of non-homolog and decoy similarity distributions

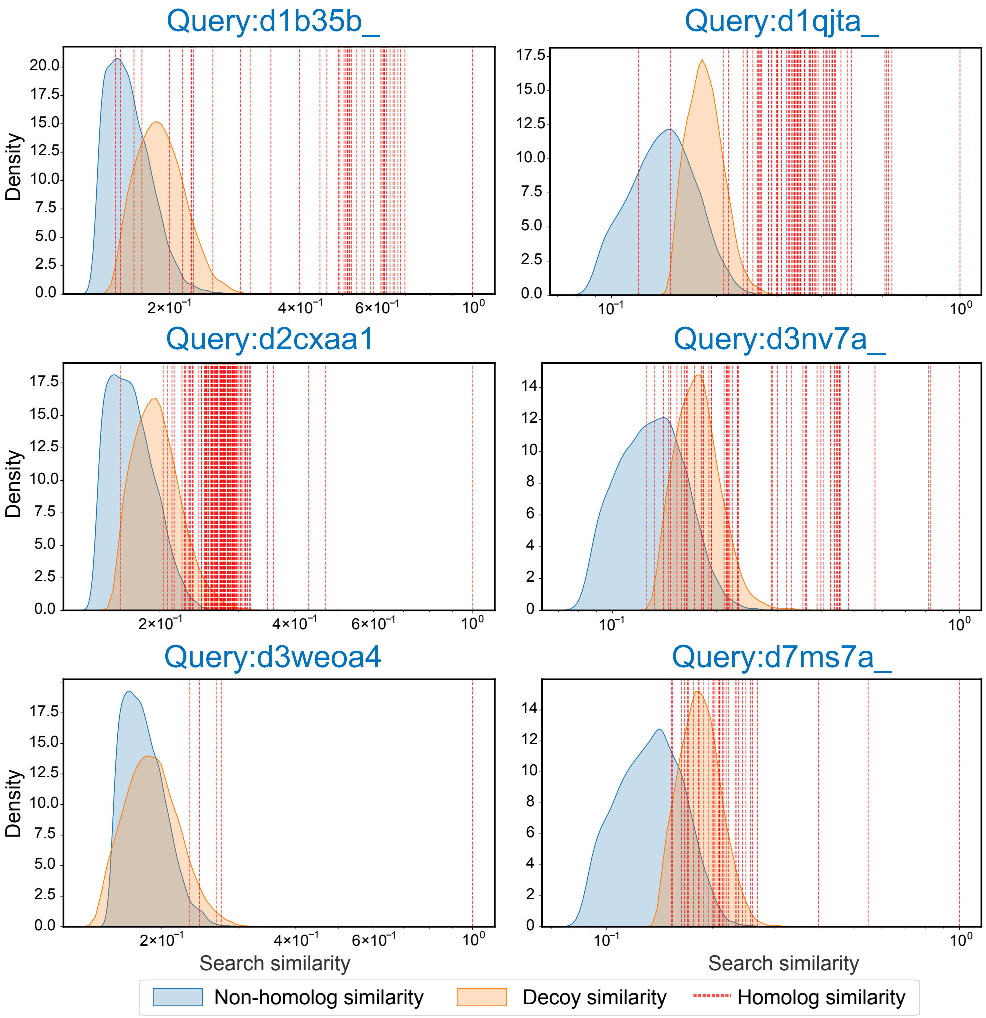

Figure S15: **Comparison of non-homolog and decoy similarity distributions across representative queries using DCTdomain.**

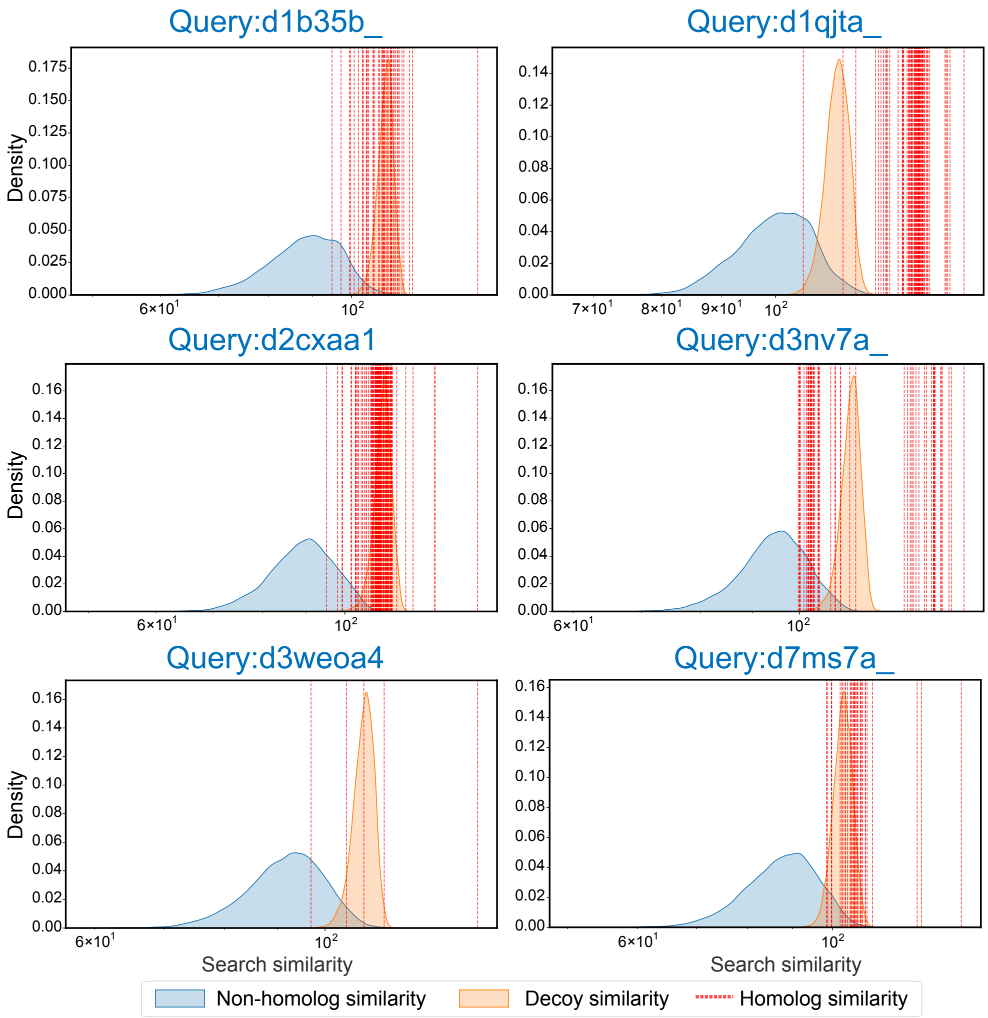

Figure S16: **Comparison of non-homolog and decoy similarity distributions across representative queries using DHR.**

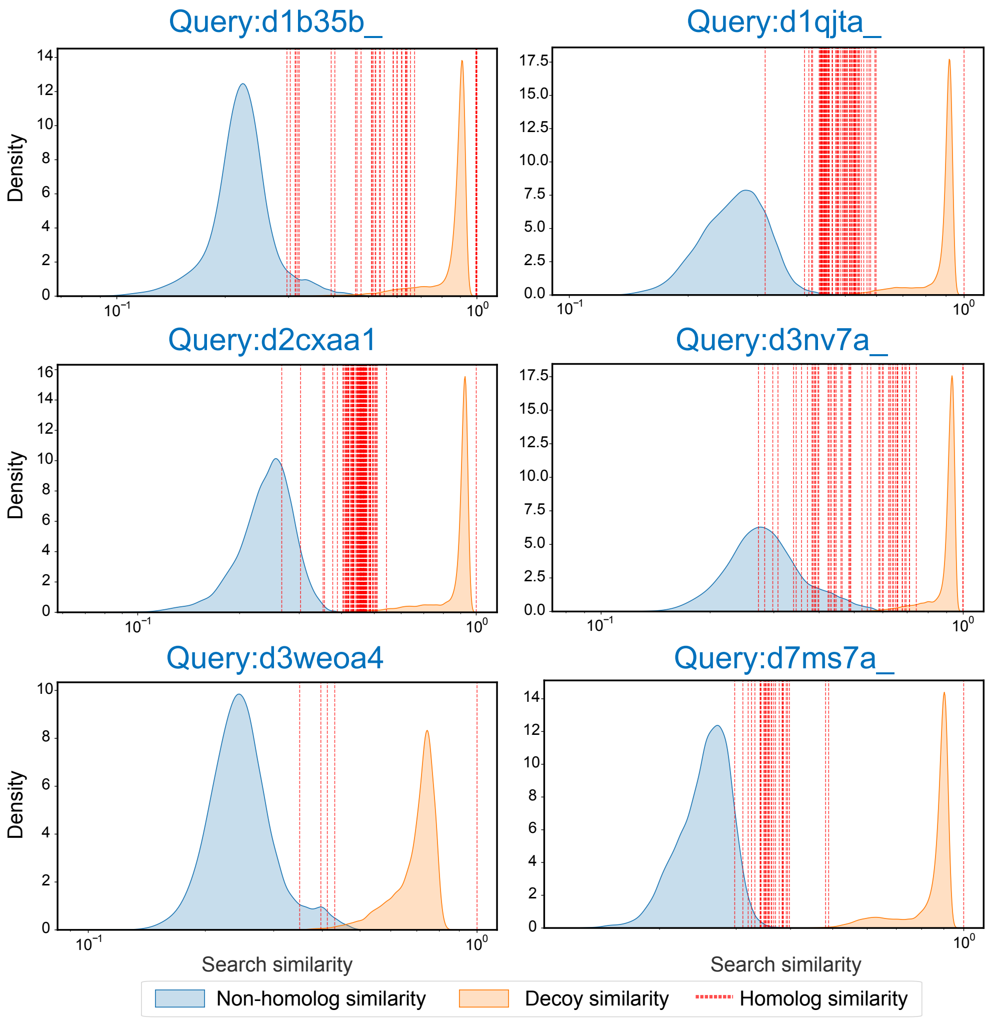

Figure S17: **Comparison of non-homolog and decoy similarity distributions across representative queries using PLMSearch.**

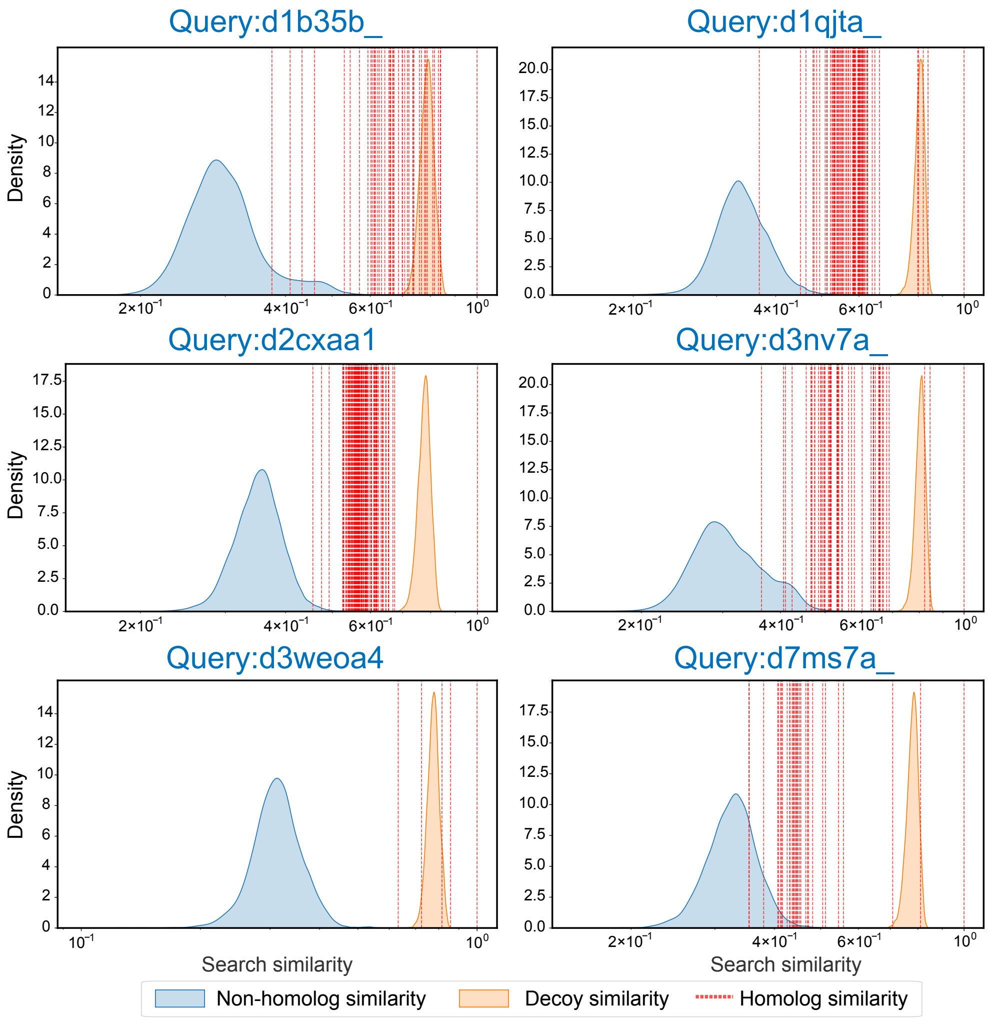

Figure S18: **Comparison of non-homolog and decoy similarity distributions across representative queries using TM-Vec.**

##### S2.3.2 Comparison of cumulative target and decoy fractions

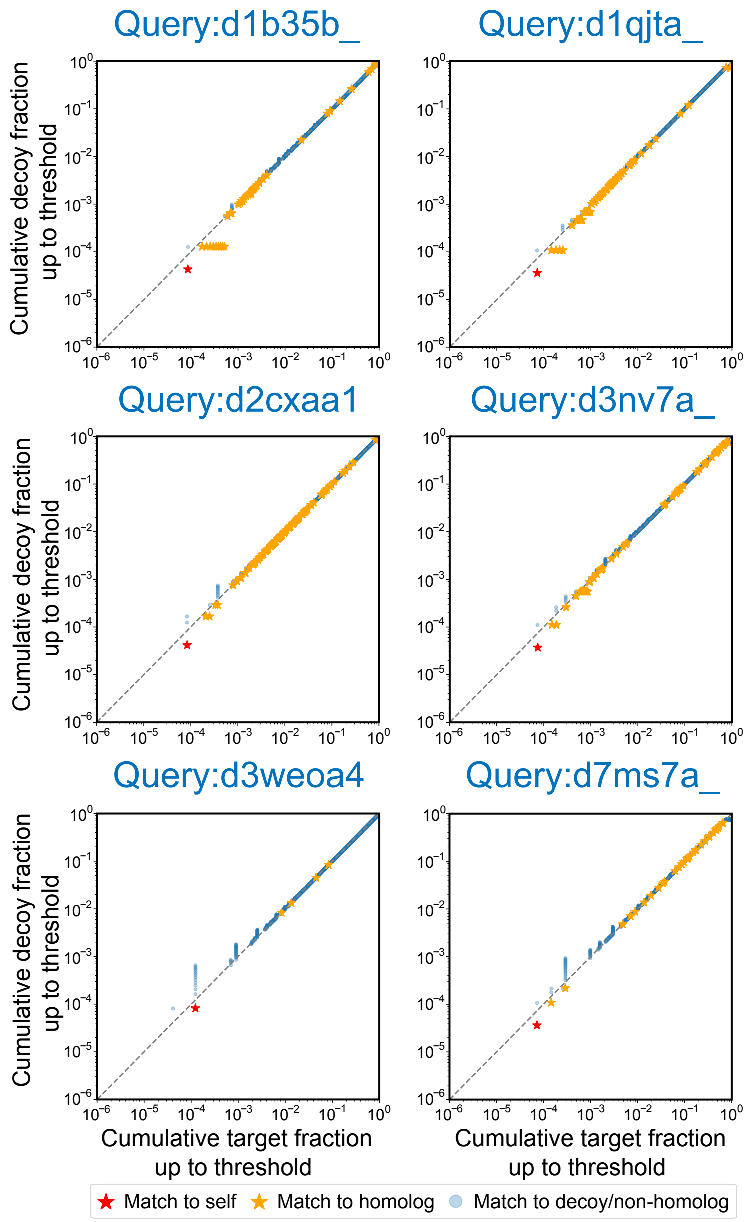

Figure S19: **Comparison of cumulative target and decoy fractions across representative queries using DCTdomain.**

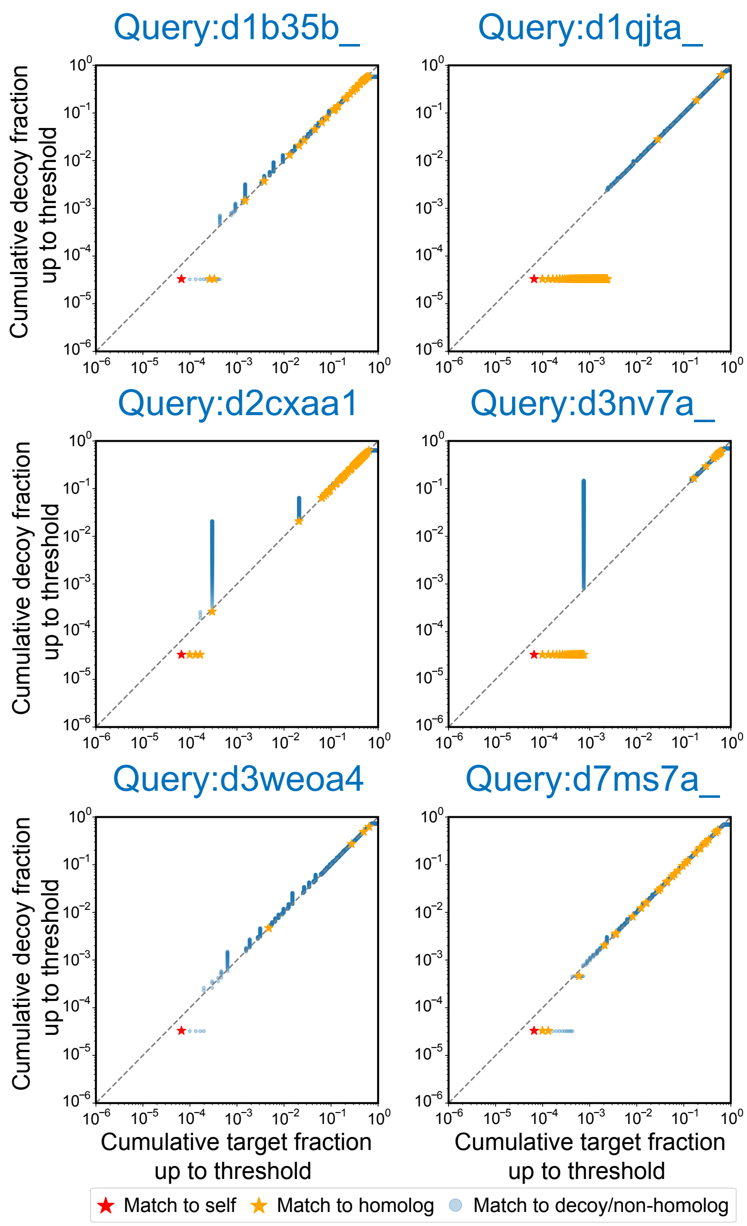

Figure S20: **Comparison of cumulative target and decoy fractions across representative queries using DHR.**

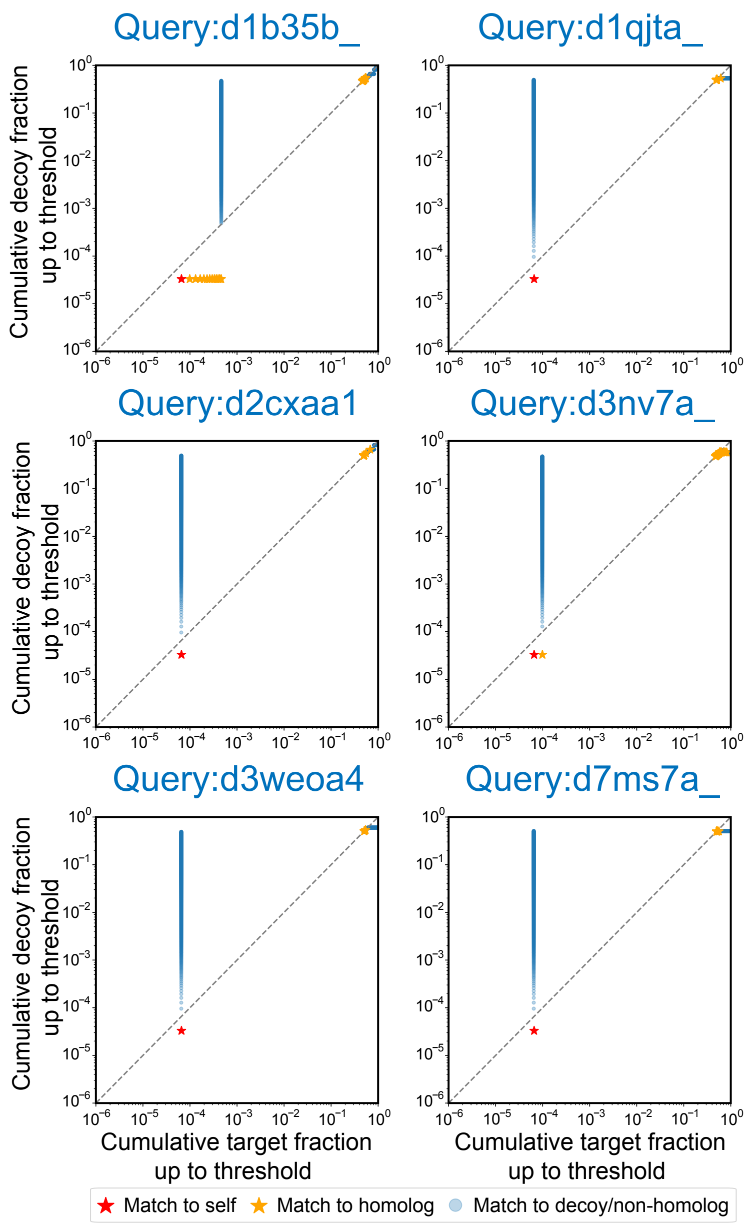

Figure S21: **Comparison of cumulative target and decoy fractions across representative queries using PLMSearch.**

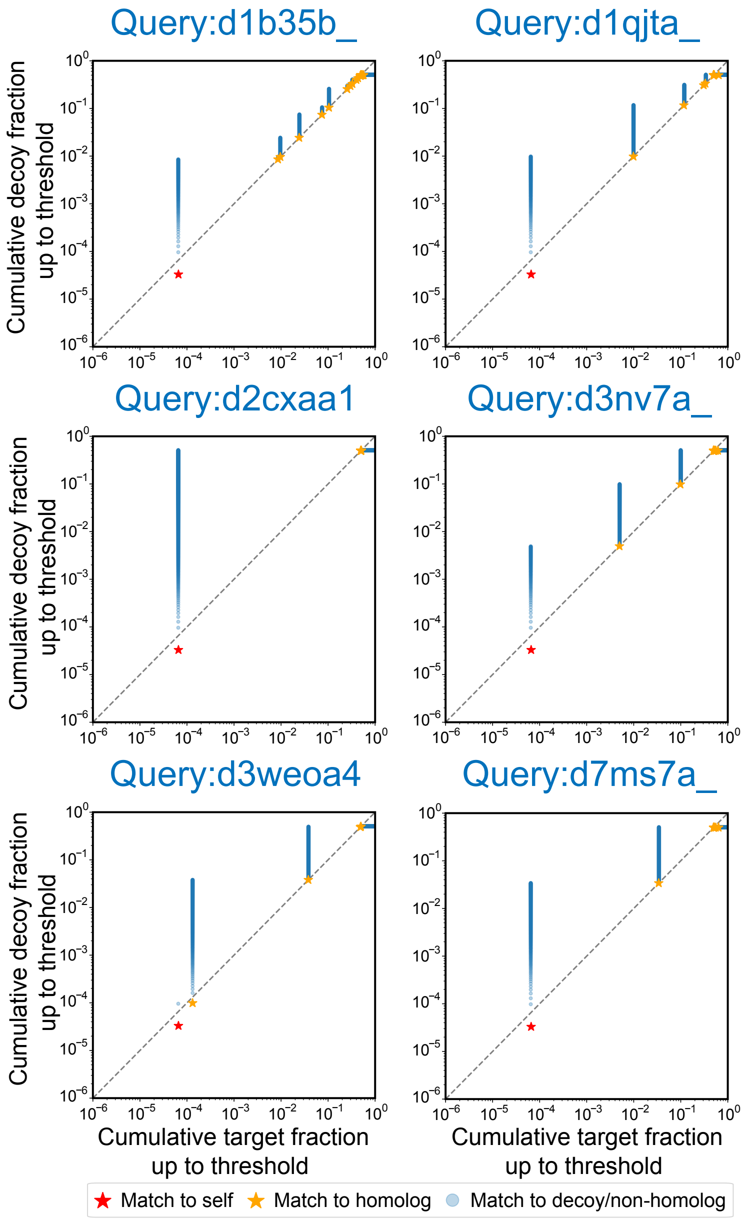

Figure S22: **Comparison of cumulative target and decoy fractions across representative queries using TM-Vec.**

### References

S. F. Altschul, W. Gish, W. Miller, E. W. Myers, and D. J. Lipman. A basic local alignment search tool. *Journal of Molecular Biology*, 215:403–410, 1990.

B. Buchfink, K. Reuter, and H. G. Drost. Sensitive protein alignments at tree-of-life scale using DIAMOND. *Nature Methods*, 18:366–368, 2021.

C. Camacho, G. Coulouris, V. Avagyan, N. Ma, J. Papadopoulos, K. Bealer, and T. L. Madden. BLAST+: architecture and applications. *BMC Bioinformatics*, 10(1):421, 2009.

T. Hamamsy, J. Morton, R. Blackwell, D. Berenberg, N. Carriero, V. Gligorijevic, C. Strauss, J. Leman,

K. Cho, and R. Bonneau. Protein remote homology detection and structural alignment using deep learning. *Nature Biotechnology*, 42(6):975–985, 2024.

L. Hong, Z. Hu, S. Sun, X. Tang, J. Wang, Q. Tan, L. Zheng, S. Wang, S. Xu, I. King, et al. Fast, sensitive detection of protein homologs using deep dense retrieval. *Nature Biotechnology*, pages 1–13, 2024.

B. G. Iovino, H. Tang, and Y. Ye. Protein domain embeddings for fast and accurate similarity search. *Genome Research*, 34(9):1434–1444, 2024.

W. Liu, Z. Wang, R. You, C. Xie, H. Wei, Y. Xiong, J. Yang, and S. Zhu. PLMSearch: Protein language model powers accurate and fast sequence search for remote homology. *Nature Communications*, 15(1): 2775, 2024.

D. Olson, T. Colligan, D. Demekas, J. W. Roddy, K. Youens-Clark, and T. J. Wheeler. NEAR: neural embeddings for amino acid relationships. *Bioinformatics*, 41(Supplement_1):i449–i457, 2025.

M. Steinegger and J. Söding. Mmseqs2 enables sensitive protein sequence searching for the analysis of massive data sets. *Nature Biotechnology*, 35(11):1026–1028, 2017.
